# Machine Learning to Summarize and Provide Context for Sleep and Eating Schedules

**DOI:** 10.1101/2020.12.31.424983

**Authors:** Tianyi Chen, Yiwen Chen, Jingyi Gao, Peiheng Gao, Jeong Hyun Moon, Jingyi Ren, Ranran Zhu, Shanshan Song, Jeanne M. Clark, Wendy Bennett, Harold Lehman, Tamas Budavari, Thomas B. Woolf

**Affiliations:** Applied Math and Statistics, Johns Hopkins University, Baltimore, MD, US; Computer Science, Johns Hopkins University, Baltimore, MD, US; Physiology, Johns Hopkins University, School of Medicine, Baltimore, MD, US; General Internal Medicine, Johns Hopkins University, School of Medicine, Baltimore, MD, US; Health Sciences Informatics, Johns Hopkins University, School of Medicine, Baltimore, MD, US

## Abstract

The relative timing of sleep and of eating within the circadian day is important for human health. Despite much data on sleep, and a growing data set for eating, there remains a need for an interpretative framework for the understanding of this data for health decisions. This study provides a new statistical and machine learning analysis of more than 500 participants in the Daily24 project. From their data, and the analysis, we propose a framework for determining the classification of participants into different chronotypes and with that the ability to realize the potential impact of daily circadian habits on health. We propose that our resulting distribution curves could be used, similar to NHANES (National Health and Nutrition Examination Survey) data for pediatric growth, as a measure for circadian misalignment and used to help guide re-entrainment schedules.

**Author summary:** Daily habits can be positive, negative or neutral for human health. Generally sleep and eating schedules are assumed without thought for their potential to help or interfere with health. In this study we propose a framework, based on data from more than 500 participants, for evaluating the relative timing of meals and sleep schedules. This evaluation, similar to pediatric growth charts, can guide clinical suggestions for those at the extremes, while helping others to realize that they are unusual relative to the population average

## Introduction

A few general rules for optimizing sleep and eating schedules have arisen from anecdotal, cultural and research based findings. For example it is now generally well accepted that eating a large meal before sleep is, on average, a poor idea for optimal health [1]. Similarly, the stress that many years of shift work places on an individual has been well documented [2]. What is not well understood is how much natural variability there is in a population of individuals with respect to their sleep and eating schedules. In a similar way, and related to the natural variability is the important, but challenging question of whether individuals can be characterized for their schedule relative to the population distribution and can be placed into different risk categories based on their sleep and eating behaviour.

For example, important milestones in an individual’s pediatric development are compared against population averages. This lets pediatricians and parents understand and even take corrective actions if the development is not proceeding normally. A similar measurement for circadian events would be ideal but is not nearly so easy to attain. For children’s measures of development, a single set of office measures and comparison against the NHANES population densities is all that is needed [3]. In contrast, for a determination of daily habits, especially ones that may have health benefits or may be dangerous to health, a set of measures needs to be performed over multiple days into the weeks or months range. Currently, complicating the comparison to NHANES, there is no similar population measure to compare the distribution of circadian measures against.

With this paper we aim to present the first steps towards the ability to measure circadian patterns within an individual and to compare those patterns against a population.

We propose to do this by building from our Daily24 data collection of more than 500 individuals who collectively contributed to an ongoing project about the timing of eating and sleeping. While this project is ongoing and still of modest size, it presents an outstanding opportunity to define what a population measure means for these types of events and how an individual can be fairly compared against that larger population.

Our framework leaves many questions open for more study. For example while we can estimate how many days an individual needs to contribute for a fair comparison, we can only do so under assumptions about the stability of a particular participants set of habits. In a related way we can posit extreme schedules for comparison to our population dataset, but we have not collected from a sufficiently large range of individuals and their behavior patterns to clearly delineate the full complexity of the measurement space. Further complicating our analysis is that there is very little data connecting long-term behavior patterns with health risk.

Despite these limitations we believe that it is important to phrase these questions and to begin the process of defining what a measure of circadian patterns should look like and how it may be used to help particular patients and their clinical care teams. We present this work with a full realization that the current framework is only a first step into this fascinating problem and we don’t believe that this is immediately ready for clinical work. In that spirit we provide an outline for how our initial dataset and analysis could be extended, validated, and eventually used in a clinical setting. We believe that efforts to establish the importance of daily awareness of eating and sleeping times can be a substantial benefit in human health and that this dimension of human health has not been fully addressed in all of its ramifications.

### Entering Assumptions and Data Collection

Participants in the Daily24 project submitted their daily eating and sleeping times through a smartphone App. We view their entries as representing stable habits that are sampled via the App on a daily basis. Clearly this is a strong assumption, since individuals may have weekly variability, may change jobs or habits, or may simply have a widely variable schedule. By collecting this information, we have the ability to define those with very regular habits, and also an ability to sense those with much wider latitude in their daily schedules.

We immediately note that we are not connecting any of the Daily24 sleeping and eating events with long-term health. This is, in part, due to the difficulty in defining the causal connections, and in part due to the limited (6-months data window) time of the study. We instead view the shared data as helping us to think about the definition of a community averaged distribution of sleeping and eating events within a (we hope) generally healthy population. We collected this data with an effort to determine daily schedules and did not instruct individuals to change their schedules as part of the study. In that regard we assume that we have a reasonable, statistically valid, distribution of collected eating and sleeping events.

The total number of lines of data was more than 1/4-million. Each of these events we view as a tweet-like note about a meal eaten, a sleep duration, or a snack. Some individuals were outstanding in contributing over the entire six-months. Other participants were less prolific in their contributions, but still did contribute a statistically meaningful set of events. We elected to trim our initial set of events to reflect individuals that contributed at least 21-days over the 6-months of the study. Each day was considered complete if it had at least a complete sleep event associated with it.

In sum the participants from this study were recruited as part of our Daily24 project and were not selected based on their particular chronotype, or on the basis of any factor known to be correlated with circadian patterns, such as shiftwork or any medical condition. We view the participating pool as reflecting generally healthy individuals, generally older, and technically literate (so that they could reliably use a smart–phone-based app and enter their eating and sleeping times).

In Fig 1 we describe the data collection process and our assumptions for analyzing the information. We make the assumption that our participants were truthful in their daily behavior and consistent in entering their schedules. We accept that we did not have the resources to verify every event and that while individual users could edit (within 48-hours) their entered events, that we did not have an ability to check each event as it is uploaded. We view this as more of a strength than a limitation, since we tried to encourage users to view this as an observational study with no intent to modify their behavior and no judgements as to the relative timing of their sleep and eating events.

**Fig 1.**
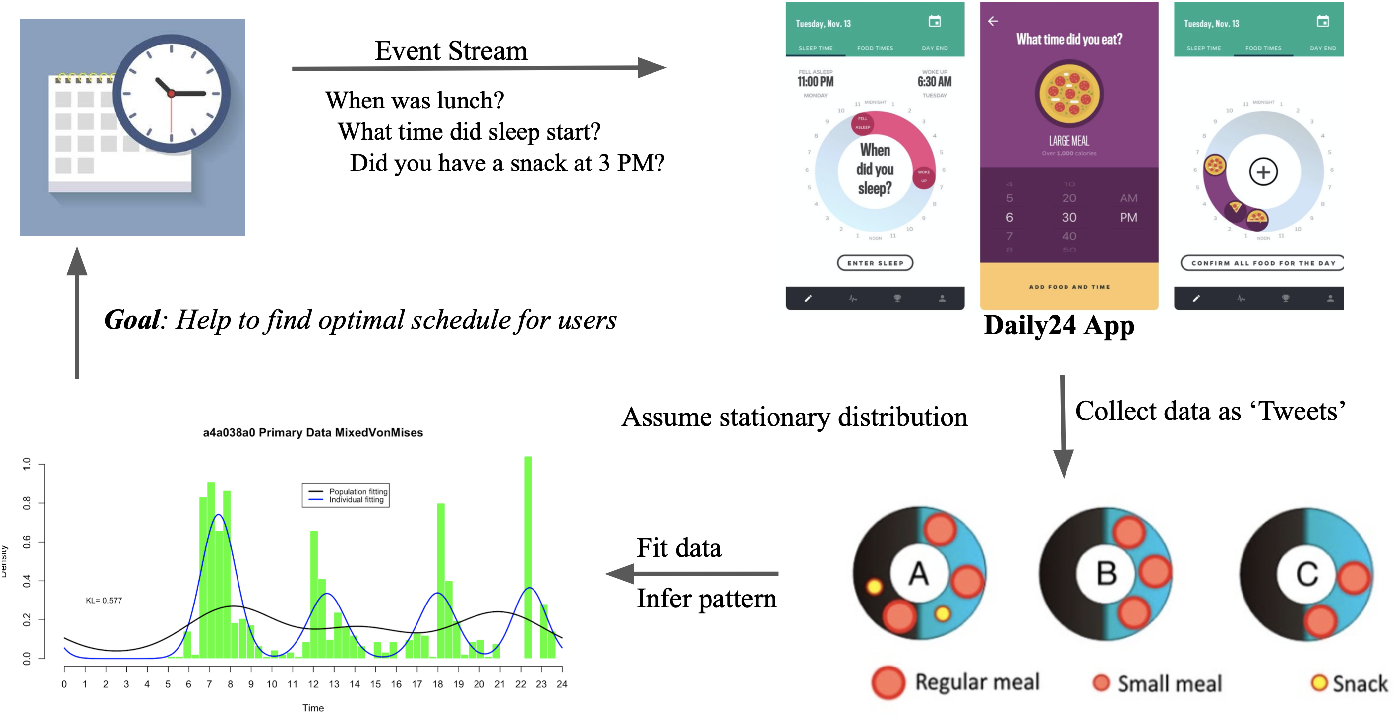
Data Collection with Daily24. Participants contributed their daily eating and sleeping habits to our Daily24 project using the associated smartphone App and our AWS backend.

With sufficient inputs, we believe that patterns in sleeping and eating can be found. As an extreme, and to illustrate this idea, it is commonly accepted that the probability of eating is not constant (a uniform distribution) over 24-hours. There are periods in the day that are more likely (greater probability) than others for an eating event to occur. Similarly, sleep is not expected to occur with a uniform probability with the most likely start of sleep occurring near the end of the circadian day.

Fig 2 shows the primary and derived data from the Daily24 App. The first set of lines is an indication of the type of primary data that we logged on the AWS backend. In each case this row included the participant’s type of entry (sleep or meal), the actual time of data-update, and some additional information. For the meal events this additional information included an estimate of meal or snack size. For sleep, the additional information included a sleep duration. We used the primary data to define a new derived data table that summarized each participant’s daily entries (one row per day). This meant that we determined a statistical fit to their meal trajectories from weak up time(calorie taken 0%) to last meal of the day(calorie taken 100%). It also meant the calculation of a meal-eating window, of a sleep window, and of the time between waking and first meal as well as of the time between last meal and sleep start.

**Fig 2.**
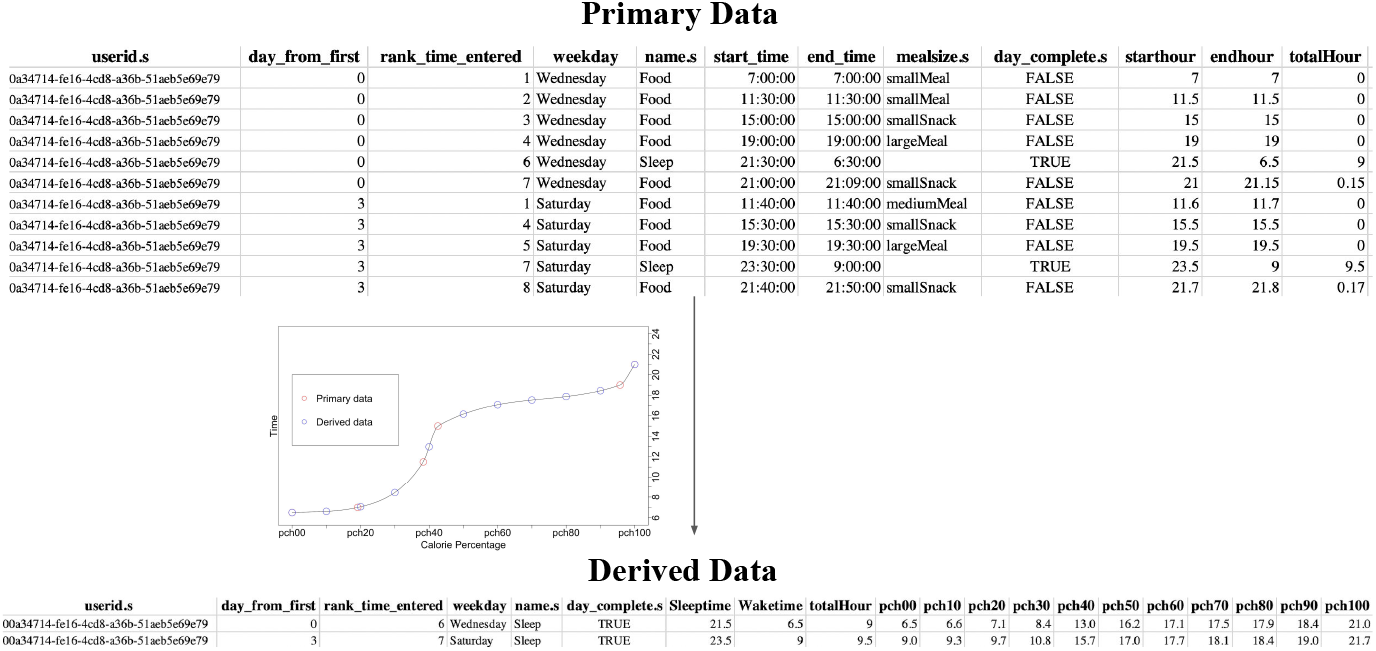
Primary and Derived Data from Daily24. Primary data consisted of tweet-like entries from the App installed on each participant’s phone to our AWS backend that stored the timing of eating and sleeping events. From this primary data the derived data summarized each day in terms of the eating trajectory and its relative length and position relative to the sleep event.

### Analytic Plan

Circular statistics were selected to account for the 24-hour cycle of our primary data. The most well-known of the distributions defined by circular statistics is called the von Mises distribution [4]. The von Mises distribution was the first circular distribution ever proposed and is arguably the most widely known and studied circular distribution. There are a few common characteristics for the von Mises distribution. It is defined by two parameters: loc, a measure of location and concentration, a measure of spread. The distribution becomes a Uniform distribution when its concentration is zero. Unlike other probability distributions, the von Mises distribution is a special case for a related distribution, the von Mises-Fisher distribution with n being two (meaning the von Mises distribution is a 2D representation of von Mises-Fisher distribution).

## Results

### Mixed von Mises

Each von Mises distribution has only one peak in its density function, while in a person’s circadian cycle, there can be several peaks. For instance, a user’s meal records may show three peaks, say at 8:00 a.m., 12:00 p.m., and 8:00 p.m., which means this user most likely had had meals at these three-time points.

Each peak needs to be represented by its own distribution, so a participant’s day of recording was assumed as a mixture of von Mises distributions, with every patient’s data represented as a sum of k component von Mises distributions, each weighted by its own coefficient. Using Bayesian framework, the conjugate prior distribution for the loc parameters was each an individual von Mises distribution; for concentration, a Gamma distribution. For the weight coefficients, we used Dirichlet distributions. We assume we know K and in practice, we tried several Ks ranging from 2 to 6 and found the one that has the best fit.

To gain an approximation of the posterior, which is the same functional form as prior distribution, we used the variational inference method [5]. Compared to sampling-based methods it has two advantages: it is deterministic and converges fast.

With this approach we gain the posterior parameters and then use the expectation of each posterior distribution as Bayes estimates for parameters in the model.

Using the mixed von Mises model, we are able to compare the posterior density functions among different users, in order to begin the process of seeing and capturing the diversity among the population. We view this distribution and approach as the simplest of those that we tried, and so its also the main reference and comparison point for the data. It is also important to reemphasize that we chose the users who were active more than 21 days as active users and took all of their records as the population. For further analysis we then compared the posterior density estimation of the population to individual densities computed for the top ten users. We calculated the KL divergence between two density functions as a numeric measure in keeping with the goodness-of-fit metric of the variational-inferential algorithm.

Fig 3 shows a fit to the entire set of individuals that contributed 21 or more days to the study data. As a comparison the kernel density estimator [6] as a fit over the histogram is also shown. By having two probability distributions we can compute the Kullback-Leibler (KL) divergence [7] between the mixed von Mises fit and the kernel density fit. The number gives us a comparison point between a well known fit to a histogram and the fit derived from our mixed von Mises model. A similar comparison by computing this KL divergence between distributions is used for providing a comparison point throughout the paper.

**Fig 3.**
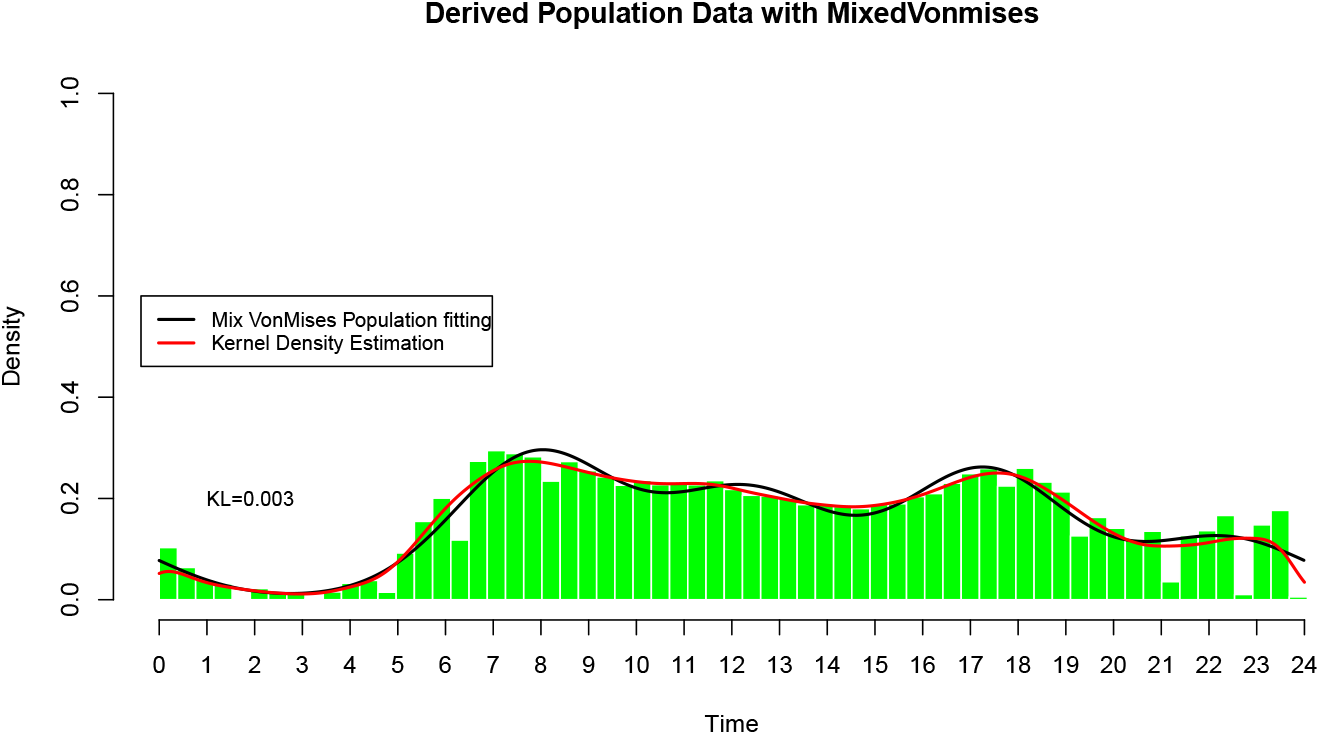
Mixed von Mises distribution fit to derived data. The Daily24 population fit with the mixture model defined by the von Mises distribution. This model provided the simplest and most consistent estimator of the densities amongst the five considered.

### Gaussian Processes

Gaussian process models have proven to be reliable estimators of many complex distributions [8]. They are a first-choice distribution for many applications in machine learning and many robust computer algorithms have been built around their application to different types of data. For our application we looked mainly at variants of Gaussian Process models that built the distributions as chains of conditional events [9]. This approach builds on the independent mixture model of the von Mises distribution by computationally arguing that the density fit is better seen as a series of conditional probability estimates. That can be simply seen as phrasing the question: given that a participant woke at 8 AM what is the probability for a first meal within one-hour?

The first of our two Gaussian process models works with code from the Turner group. In particular, their GPAR stands for the Gaussian Process Autoregressive Regression Model [9]. GPAR models are multi-output regression models used to exploit dependencies between outputs to maximize predictive performance where it can also capture nonlinear relationships between outputs. One specific feature that we thought GPAR useful for is its ability to define functions in terms of each other and to then stack them together.

In order for GPAR to work for our data, we first ordered our data into three separate data sets: wake time, food time, and sleep time. Then, we ordered the food times by ordered meals: first meal, second meal…, last meal. For most top contributors, the total meal numbers ranged from six to eleven meals. After we ordered the data in the following way, we defined distributions for each event. When we fit the Von Mises distribution, there were many meals that overlapped with each other. As a result, we have combined a few meals together for the individuals that were close to each other so that there will be four meal events.

After working through the data component of GPAR, we defined each function to be von Mises distribution. Because every event should start with the wake-up time, we have used the same von Mises distribution fit defined previously. On the other hand, for other events, we created sample points from previous events and refitted so that we have functions that depend on both the data and the previous events. Afterward, we stacked them and added some noise. Now using GPAR’s regression function GPARRegressor, we fit our functions to the Gaussian Process model. Then, we plotted observed points, the von Mises line, and the Gaussian Process Regression.

Multi-Output Gaussian Process Toolkit (MOGPTK), our second Gaussian Process approach builds from a Python package for multi-channel data modeling using Gaussian processes (GP) [10,11]. This toolkit aims to address the need for a Multi-output Gaussian Process kernel and provides a natural way to train our model and it is based on the trained model to predict the following pattern. To apply this toolkit, we also need to implement GPFlow [12], which is an extensive GP framework with a wide variety of implemented kernels, likelihoods, and training strategies. MOGPTK is a based MOGP kernel from which specific kernels are generated. The base kernel provides the functionality to split the input data into multiple channels and process them by sub-kernels.

Under this toolkit, we mainly focused on the MOSM kernel, which is the Multi-Output Spectral Mixture Kernel [13]. The MOSM kernel is designed to provide a closed-form covariance function after applying the inverse Fourier transform. Based on the Parra and Tobar paper, the cross-spectral density between channels i and j is modeled as a complex-valued SE function. In our approach, we applied the MOSM Kernel to find the covariance, mean, magnitude, delay, and phase between every two variables, and then built our model under the MOSM Kernel.

Fig 4 shows the fits from our two Gaussian process models and the KL divergence relative to the kernel density estimators. Note that both models give good fits and that this enables a comparison to the mixed von Mises as well as between the two GP models.

**Fig 4.**
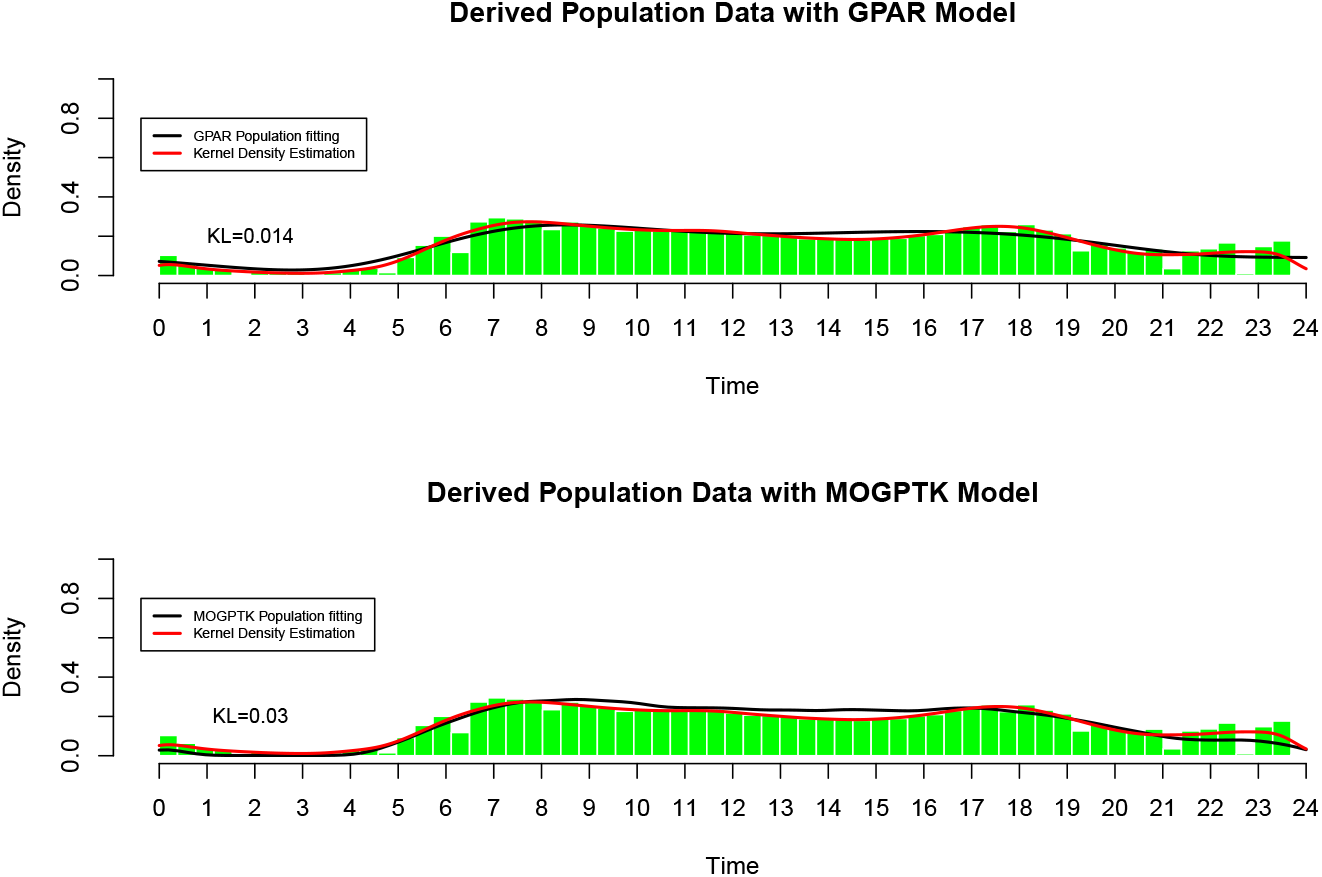
Gaussian Process Models. GPAR and MOGPTK models were defined for population fits to the derived data of the Daily24 population. Note the comparison back to the mixed von Mises and to the next figure for the State Space Models

### State Space Models

The state-space model (SSM) is a statistical model that relates a set of observable variables (so-called manifest variables) to a set of latent variables [14, 15]. It is assumed that the responses on the indicators or manifest variables are the result of an individual’s position on the latent variable(s), and the manifest variables have nothing in common after controlling for the latent variable. We used SSM which is a package in Python consisting of fast and flexible code for simulating, learning, and performing inference in a variety of state-space models.

In order to make state-space model work for our data, we converted our data to “counts” type for 8 behaviors as realization of observable variables (day-count for ‘wake’, ‘drinkOnly’, ‘smallSnack’, ‘largeSnack’, ‘smallMeal’, ‘mediumMeal’, ‘largeMeal’, and ‘sleep’). This reflects the type of data that the SSM package is expecting to use for fits. Our aim is to estimate the underlying latent variables, which in our case can be interpreted as the individual’s energetic level, see Fig 6, 7, 8.

Specifically, we used hidden Markov model(HMM) [16] and switching linear dynamics system(SLDS) model [17] to fit the data. HMM as a classic state space model is simple and easy to fit while SLDS can interpret data more subtle with continuous hidden states [18].

To compare the two state space models with the GP models and the mixed von Mises, Fig 5 gives a comparison. Again, the kernel density estimator is used to provide a comparison point.

**Fig 5.**
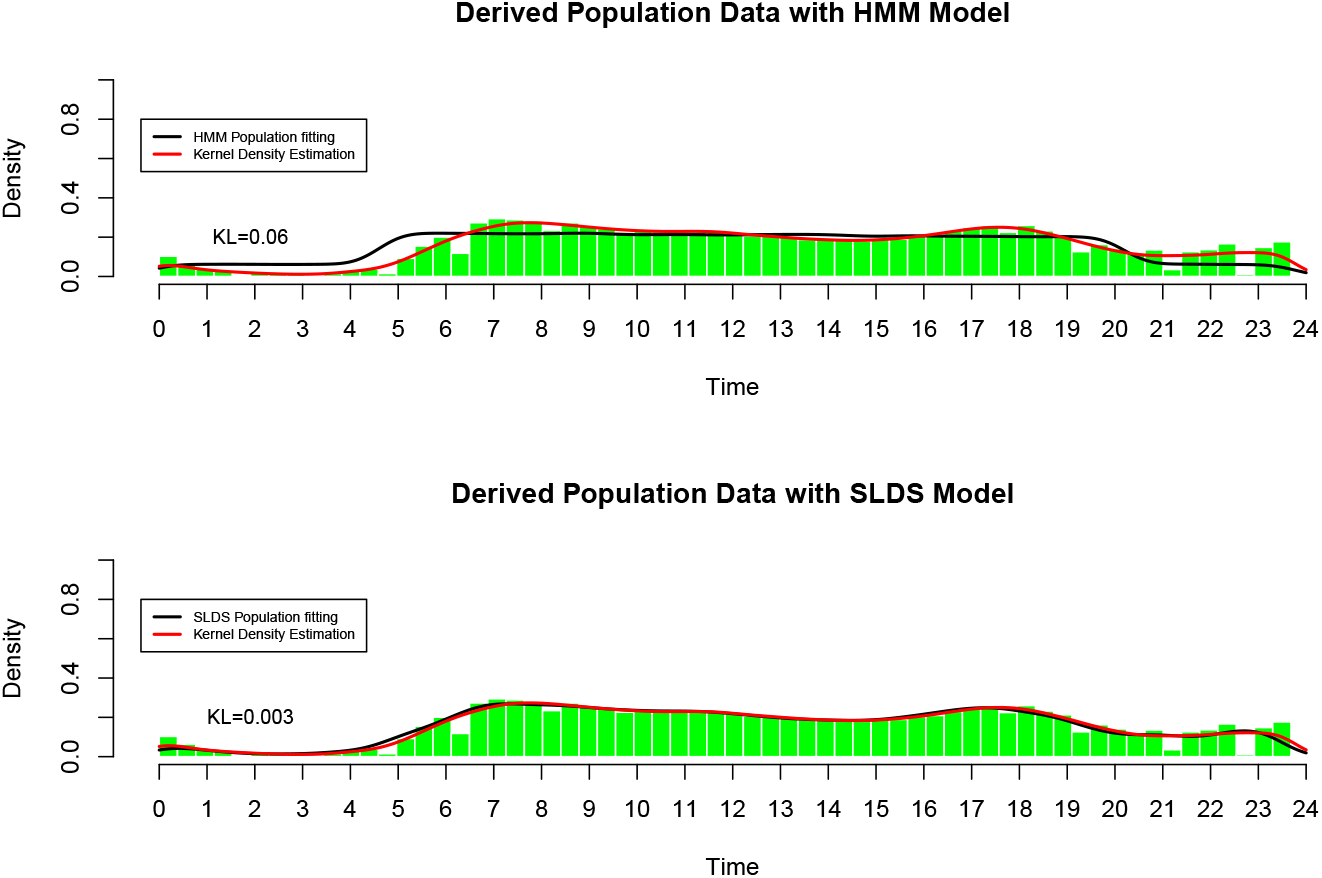
State Space Models. HMM and SLDS models were defined for population fits to the Daily24 data.

We conclude this presentation of results for the population fits by reference to Table 1, where the different models are compared. Note that while there are differences between the models, these may not be sufficiently different to enable a clear winner or subset of losers in the models to be discriminated. For that reason we evaluated how individuals are scored in the following section.

**Table 1.**
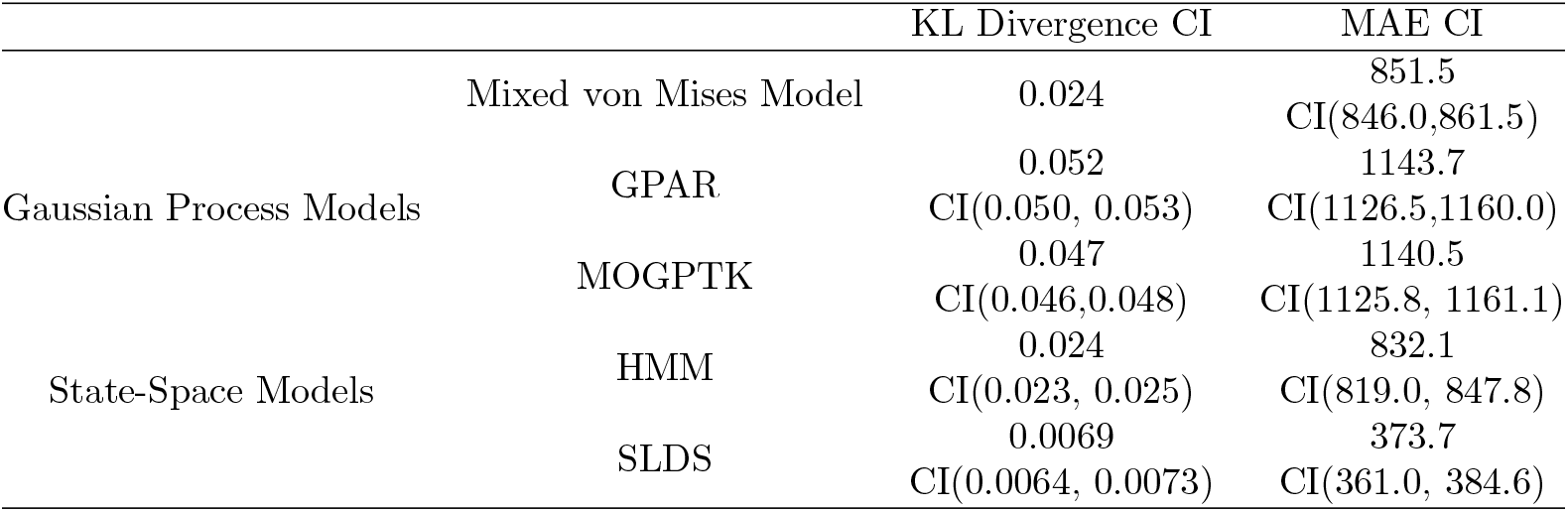
Comparison summary table for 5 methods on primary data. We use 2 error metrics: KL divergence(KLD) and mean absolute error(MAE). We calculate KLD between each method fitting and Gaussian kernel density estimation(Gkde) of the data.(Since *KL*(*p,q*) ≠ *KL*(*q,p*) we use mean of two values) For mixed von Mises model, the calculation is straightforward, for GP models and SSM models, we can only sample from model results and get data counts for time windows. By this way we recover a sample data(size 1000) from each model and apply Gkde on the sample data, compare to Gkde of original primary data, then get KLD. Replicating this procedure for 100 times gives estimation of KLD and corresponding confidence interval(CI). For MAE, we compare sample data from each model and original primary data with histograms(48 bins), then calculate mean absolute difference of counts in each bin as MAE. GP models and SSM models’ MAE calculation is straightforward, we use inverse function method to sample from estimated mixed von Mises distribution. Replicating this procedure for 100 times gives estimation of MAE and corresponding confidence interval

### Comparison of Individuals to Populations

To evaluate our five different models we explored how individuals were scored in each of the five models. This did not lead to a clear winner (i.e. one best model), but did let us evaluate how different types of individuals would be represented within the different formulations.

What we feel is thus important, for model selection in this instance, is to closely evaluate what properties of which individuals are most critical for a clinical application. Since we do not have that additional data there is not easy way to discriminate between the different models.

Instead we are left with a set of differences that are interesting, and are presented here for comparison. For example, in Fig 6 the way an early activity individual would be seen in the five models is shown for comparison.

**Fig 6.**
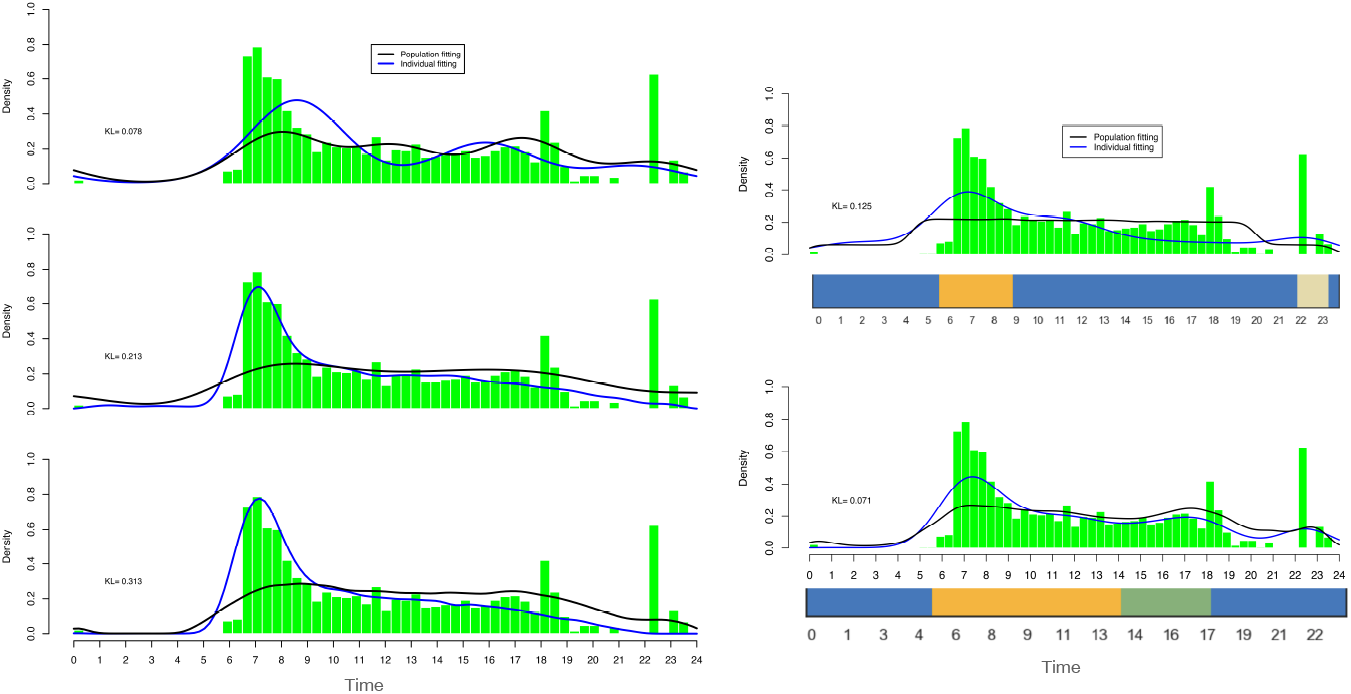
Early activity peaking individual. Comparison of all five models for one individual (a4a038a0). Note the variations in KL score. HMM provides three states where the orange one can be viewed as the most energetic period during the whole day since the individual begins working in an early time, then the change to an ivory state in the night may reveal a transfer to rest time and blue region may stand for the whole other time. SLDS method also gives three states which we may also interpret as the energy transferring, up to the difference brought by the duration of each period.

Fig 7 shows a similar comparison figure for a late peaking individual. Note for both this figure and Fig 6 that the state space models have a slight advantage in providing a sense for what hidden degrees of freedom might be involved with setting the observed distributions. The quality of the fits are sufficiently similar in all five models for both early and late active individuals to be sufficient for our goals.

**Fig 7.**
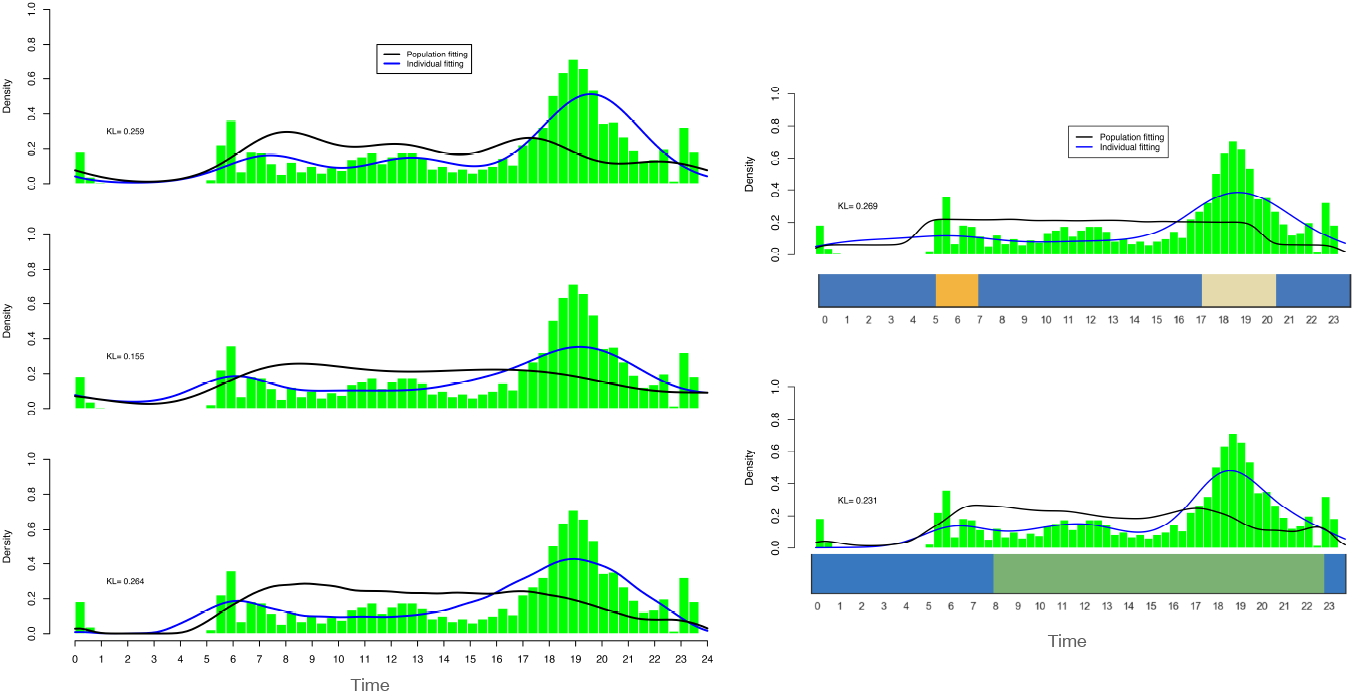
Late activity peaking individual. Comparison of all five models for one individual (e6e4ca7f). Note the variations in KL score. HMM gives three latent states, the ivory one may display the working time, the most energetic period, the orange one may indicate the waking time while the blue one represents the rest time of the whole day. SLDS gives two separate states, the blackish green can be viewed as the whole period during which individual takes most of his activities while the blue one may represent his rest time.

In Fig 8 we illustrate a challenge for all five models: how to capture an individual with both late and early peaks in activity. In this case the mixed von Mises and the two state space models provide better fits. Note that only one of the two state space models (HMM) is able to give some indicators of internal variables that underlie shifts differing between early and late initiations.

**Fig 8.**
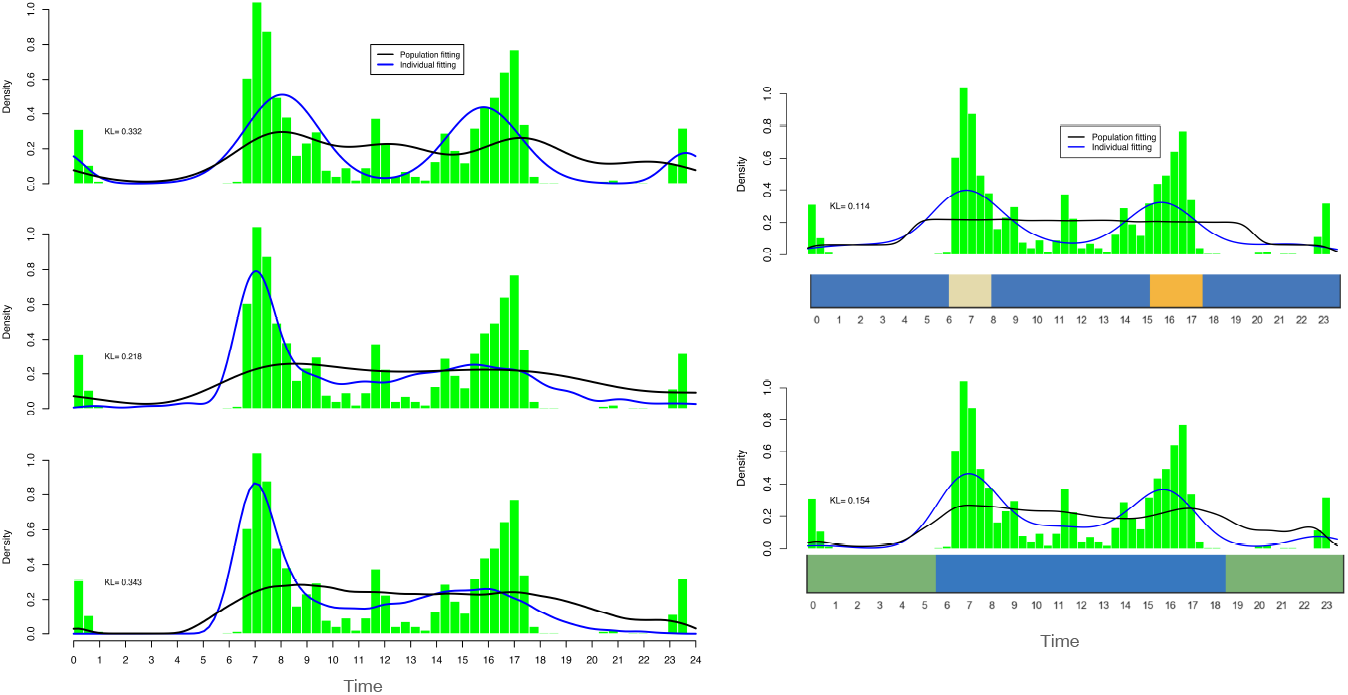
Early and Late peaks in activity. Comparison of all five models for one individual (448a8708). Note the variations in KL score. HMM provides three latent states. The ivory one and orange one are two periods corresponding to the two peaks in activity and the blue one manifests the other time. SLDs offers two states where the blue one exhibits a longer duration which may be interpreted as the whole time period where the individual conducts his activity and the blackish green one signifies the rest time.

One way to further compare the different models is to plot the individual KL divergence scores for each model relative to a common reference probability. This is shown in two ways in Fig 9 and Fig 10. Note that a set of individuals (8 in total, three of them are shown in Fig 6, 7, 8, while the rest are in the supporting information) for comparison is shown along the x-axis. For Fig 9 the ordering along the x-axis is from most consistent score to most divergent score. The distribution of scores in Fig 10 illustrates that there is a non-symmetric distribution with a very long tail on the right hand side.

**Fig 9.**
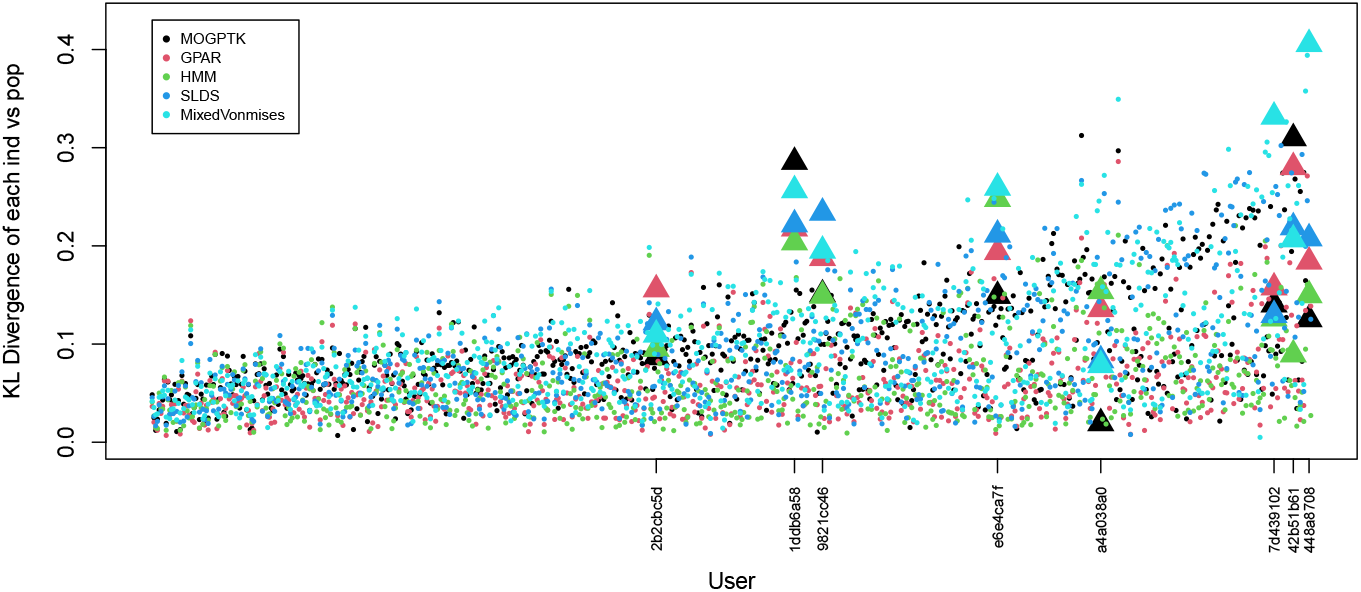
KL divergence as scatter plot for individuals in each of the five models. The diversity of measures shows that non-uniqueness of the fit with some models measuring individual differences as more or less extreme relative to their estimate of the population distribution.

**Fig 10.**
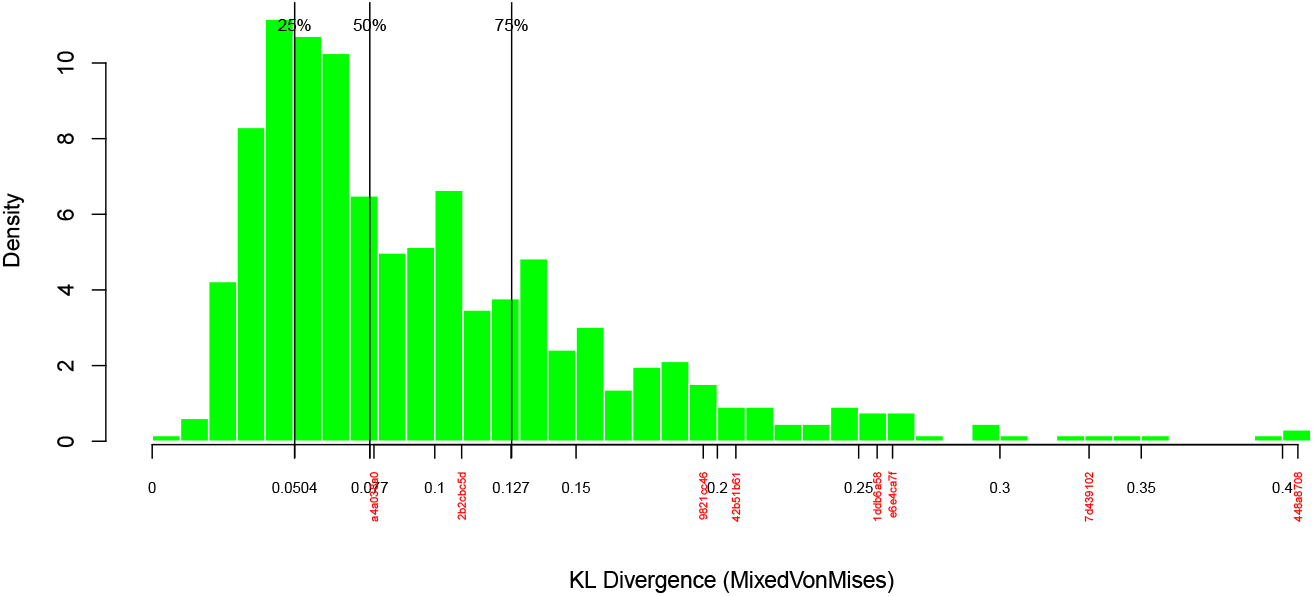
Histogram of KL divergence within the von Mises fit. The diversity of measures shows that non-uniqueness of the fit with some models measuring individual differences as more or less extreme relative to their estimate of the population distribution.

## Discussion

Circadian biology is an ancient part of human physiology and reflects our human evolution within the context of a twenty-four hour day [19]. The adaptations to the light/dark cycle of each 24-hour cycle has been an important regulator of many biological systems. While the molecular, cellular, and tissue ramifications of these adaptations remain an active area of research, every individual makes decisions about eating and sleeping without much thought or context on each day.

Recent work has shown that the timing of eating has a major impact on circadian function [20]. The beginnings of circadian physiology emphasized light and sleep as the main drivers of circadian rhythms. With the realization of the importance of the timing of eating, the full awareness of the coupling between peripheral and central components of circadian biology came into focus. With this awareness has come an improved ability to define circadian mis-alignment as due to behavior (for example shiftwork) that does not support a consistent 24-hour rhythm that aligns light/dark, sleep and meals [2].

With the growing acceleration of technology the ability for many individuals to ignore light/dark and to work at many hours has become common. While the new habits that this brings may seem to have only trivial impact on a daily level, they can lead to significant physiological stress over years. To evaluate the relative impact of the daily habits over many years is a challenge that is not yet fully addressed. To help interpret what a circadian daily habit means for human health there is the need to a summary of many days of behavior and a way to relate the individuals behavior back to both optimal behavior and to the statistical behavior of many others. The NHANES project has provided many families and pediatricians with a dataset that lets a comparison of an individual child relative to the population to be easily defined [3]. Our work is in the same tradition, with the expectation that the methods defined by this paper an provide the entry point to a larger, community defined, dataset for summarizing, interpreting, and aiding, in the evaluation of circadian health.

### Changepoint Detection

It has been observed that many individuals have different weekday versus weekend schedules. Often this is modeled ‘as if’ the individual has shifted timezones. This framework should also be readily interpretable for the Daily24 data. But, we did initially fit all of the data ‘as if’ each day is identically distributed. Our entering assumption thus made the model fit easier, but at the possible expense of leaving out the true complexity of individual behaviour.

We address this issue by bringing in our initial efforts at changepoint detection. This is shown in Fig 11 where we demonstrate that it is possible to optimize a scoring parameter for our data. In principle it should be possible to further determine by the sampled Daily24 events whether someone has an unusual schedule (for example Wednesday is always different from all other weekdays) or whether a particular Wednesday is simply different from other Wednesdays. This is a challenging problem with many algorithms that have been defined for changepoint detection, but without a clear indication for when a particular data point represents a genuine change or simply a fluctuation within the same stable distribution.

**Fig 11.**
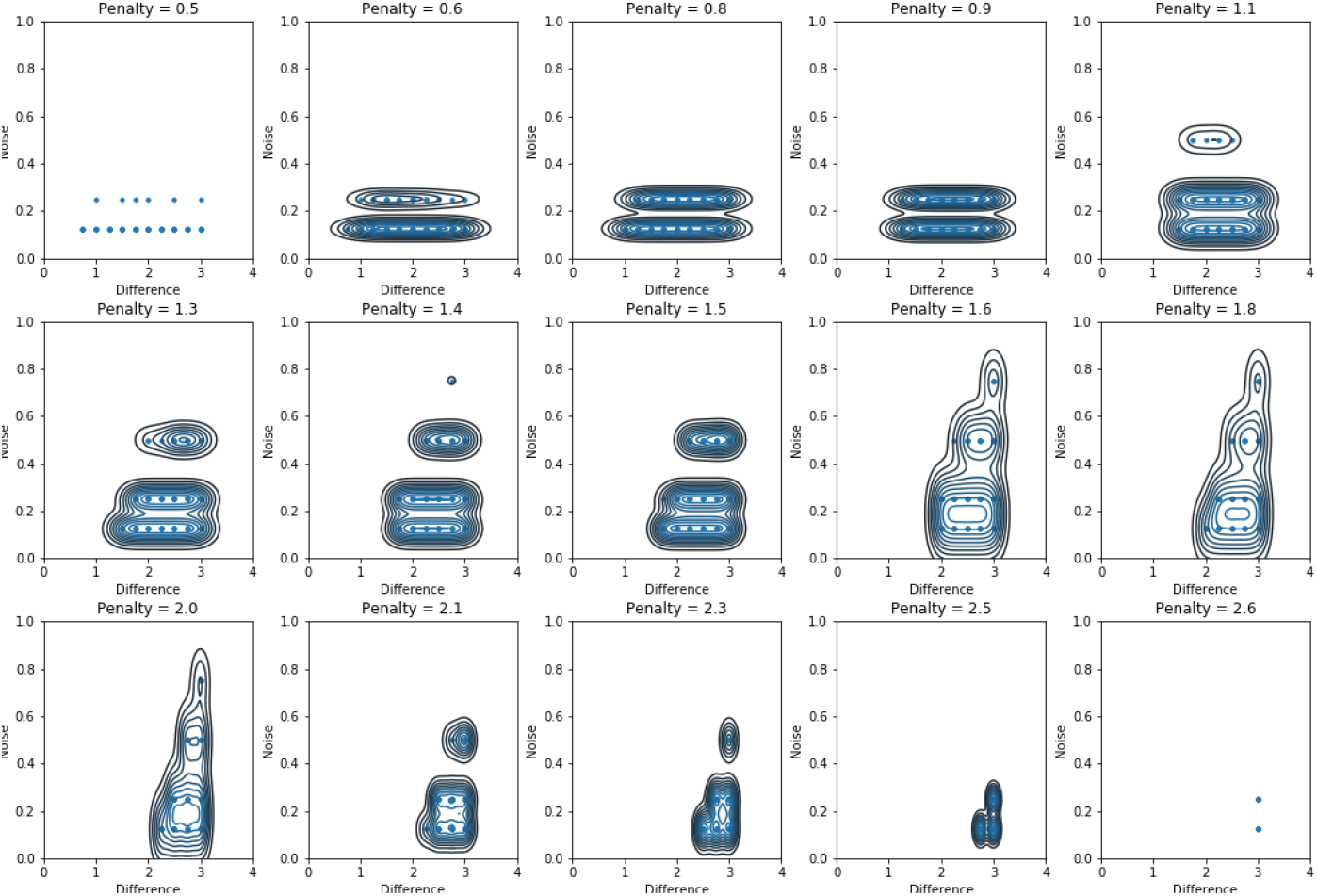
Defining the optimal penalty function for the Changepoint Detector. Participants will change their schedule, sometimes due to a work change, sometimes due to a weekday versus a weekend. To simplify the initial analysis, we made the very strong assumption of a single stable pattern. To define when a shift from that pattern is due to a schedule change versus an unusual event from the same stable distribution is the challenge of optimizing a changepoint detector. We present, in this figure, the first steps into defining one for Daily24 data.

An additional type of changepoint is that seen with shiftwork. While we did not have participants in Daily24 that shared a schedule similar to the traditional shiftwork schedule, we do feel that the Daily24 App should be applicable to shiftwork schedules as readily as the ‘normal’ schedules that we sampled. In addition, the changepoint algorithm should be capable of detecting shifts in daily schedule that reflect an ‘on-shift’ day relative to days that are ‘off-shift’.

The changepoint approach that we used is built within the Rupture Python code and is based on multiple papers [21]. We evaluated a range of different possible algorithms and different penalty functions. While we were not able to tune the changepoint detection to reliably get all changes correctly labeled, we did get results that suggest the implementation of a changepoint detection would be important for the generalization of our results. In particular, we suggest that changepoint measures, even if imperfect, can be a large help in identifying those individuals with a large weekend effect or that have shiftwork schedules.

### Cluster Analysis

To understand whether different daily habits can be classified we created a set of synthetic individuals with very regular schedules. This let us supplement out Daily24 samples with individuals that represent early and late chronotypes, with shift workers, and with those that consistently eat meals before sleep. For more details on the synthetic datasets please see our supplemental information.

This supplemental dataset lets us define a feature set that includes both a comparison of each individual to population measures using KL divergence and an analysis of changepoints in an imagined daily schedule. By using this feature set we can readily classify individuals into different types. For example the early and late chronotypes are clearly discriminated from the other types of behavior in this classification. This is shown in Fig 12

**Fig 12.**
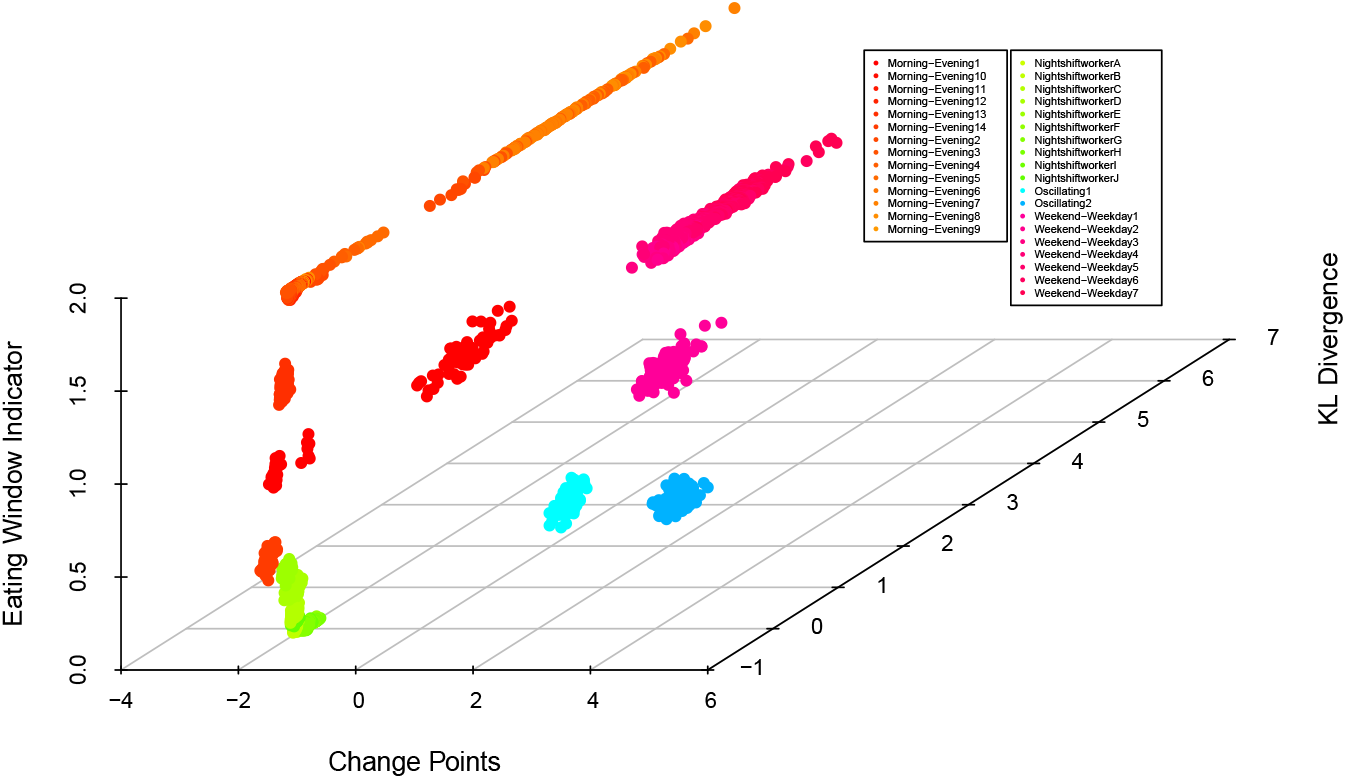
Classification of an individual based on timing of sleep and of eating. Synthetic data is used to illustrate how a machine learning framework for classification would enable confident mapping of individual behavior to a population based understanding of relative risk and of patient type.

### Clinical Presentation

While the analysis we present is not ready for clinical work, the approach that we outline may provide a framework for how daily habits and their summary can be presented within a clinical setting.

As an example we imagine that the synthetic data is a reasonable representation of the range of observed human behavior. This let us describe the approach, but we emphasize that without real participant data that the ideas presented are still at the idea stage.

With the assumption of five converged population distributions, representing the early and late chronotypes, the shiftworkers, the weekend/weekday shifting schedules and those with a long-term habit of late meals, we can define each person’s circadian schedule relative to their main reference population. We can imagine a type of presentation that is summarized in Fig 13 where each individuals set of values is compared against the population measured distribution. This is a similar spirit to the NHANES data for pediatric growth and could serve a similar purpose by providing a discussion point between clinical providers and their patients.

**Fig 13.**
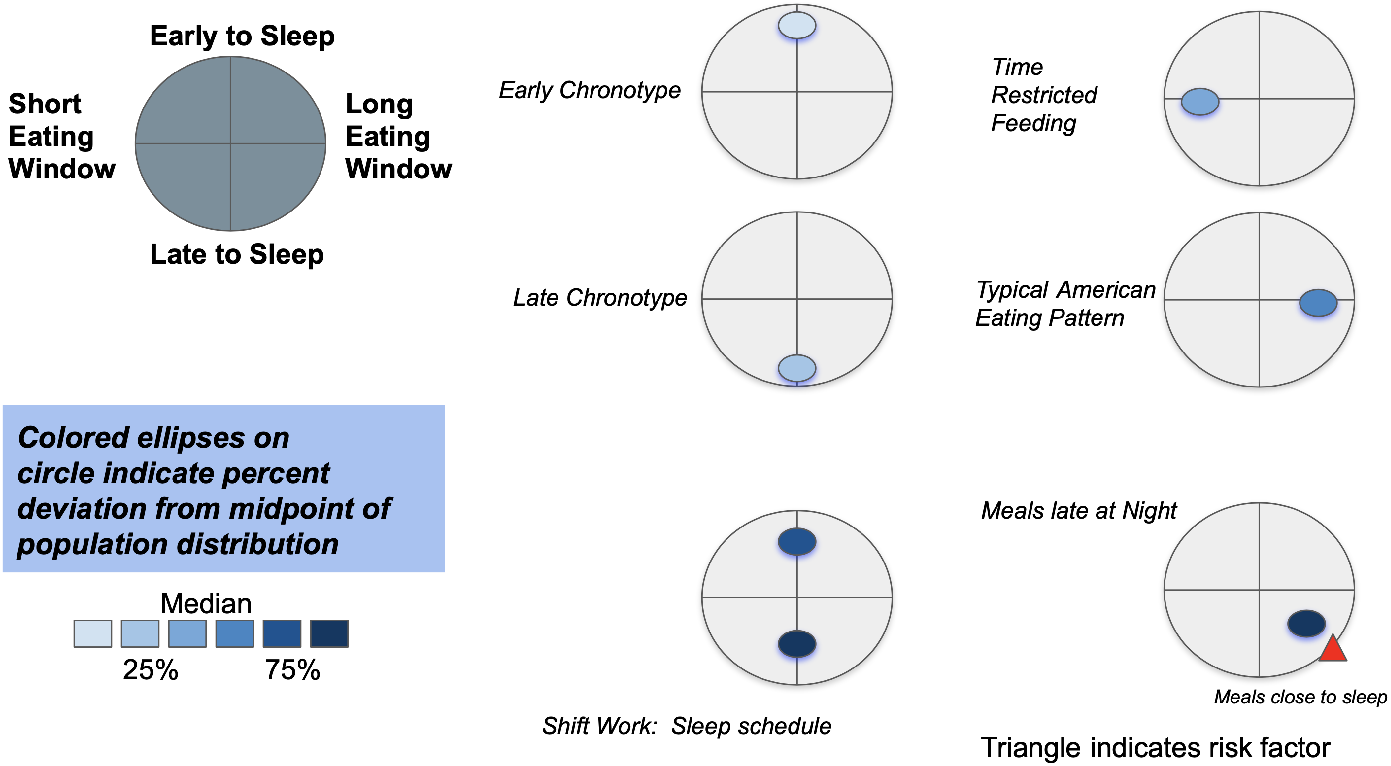
Interpretation of data with chronotypes and for the clinical setting. To enable clinical care teams and patients to readily see what their data means and to help with communication, something like this data presentation may be used. What is plotted is the strongest set of events distributed over the participants logged timings for eating and for sleep. By plotting this on a relative axis, with additional use of color and symbols, the clinical meaning can be made clear.

This may be especially important for clinical issues surrounding circadian mis-alignment, for example in high-risk populations like shift workers [2]. We can imagine the ideas of this paper being combined with ideas for optimal re-entrainment from travel with light schedules to define new schedules for optimal recovery from mis-aligned light/dark and eating/sleeping [22, 23].

An additional focus area for this type of analysis is for providing a continued assessment of the impact of intermittent fasting on human health. This was our own entry point to Daily24 and the analysis of this paper could be used to more fully characterize the impact of timing of eating by a comparison back from individuals to the community. There has already been much valuable research on the impact of timing of eating, and this framework could help to move these types of questions from individual anecdotal studies or from purely research based studies, into a clinical setting by providing a consistent framework for comparison [19, 20]

## Conclusion

A significant unsolved problem for circadian physiology is how to extrapolate from a daily habit into the impact of that habit over years. While our current study doesn’t connect the health outcomes with the daily habit, it is a framework for providing an ability to quickly summarize the daily habits of an individual within the context of a larger group. While the direct clinical application of this approach will need still larger datasets and still more analysis work to connect the distributions to health outcomes, we believe that this framework approach can provide an important anchoring point within a population for the interpretation of many days of circadian data. This is a real improvement over a scatter chart of an individual’s data and potentially can provide a way for nuanced discussion of an individual’s daily habits within the context of a clinical office visit.

We emphasize that the approach outlined will need to be extended to a larger dataset with more varied individuals (by age, gender and race). The importance of potential cultural bias (since all participants are in the US) is also a factor that should be considered in enlarging the dataset.

Furthermore, the optimal approach for how long an individual’s circadian schedule should be tracked and with what confidence bars will need to be worked out. While some confidence bars can be supplied based on assuming that (for example) a two week schedule is fully representative of a two-month or two-year schedule, this may clearly lead to a large systematic error if the assumption of an unbiased and consistent sample is wrong.

## Supporting information

**S1 Individual example-1.** Active more in the early part of the day.

**S2 Individual example-2.** Active throughout the day.

**S3 Individual example-3.** Active later than most.

**S4 Individual example-4.** Active more in the early part of the day.

**S5 Population fitting on primary data**

**S6 Github Repo** contains codes for all five methods and an synthetic data for each method to run. See https://github.com/jdjmoon/TRF for details

## Ethics Statement

Our Daily24 app data was abstracted from our larger AHA study. This larger AHA study was approved by our Johns Hopkins IRB panel (number: IRB00174516). All participants in the study signed an informed consent form. Additional data, not included in this paper’s analysis, included REDCAP survey questions and selected EHR data all approved by our IRB and collected with informed consent.

## Acknowledgments

This study was made possible by the AHA funding for a Strategically Focused Research Network (SFRN). We want to thank the participants, staff, and research skills of those that helped with the Daily24 project. We also benefited from the AMS Department (Homewood) in the start of the analysis for this project, the use of IDIES resources (SciServer) and want to thank other members of the class and departments.

