## Supplemental Material for "Machine Learning to Summarize and Provide Context for Sleep and Eating Schedules"

### Individual fitting on derived data

#### 1ddb6a58 Active more in the early part of the day

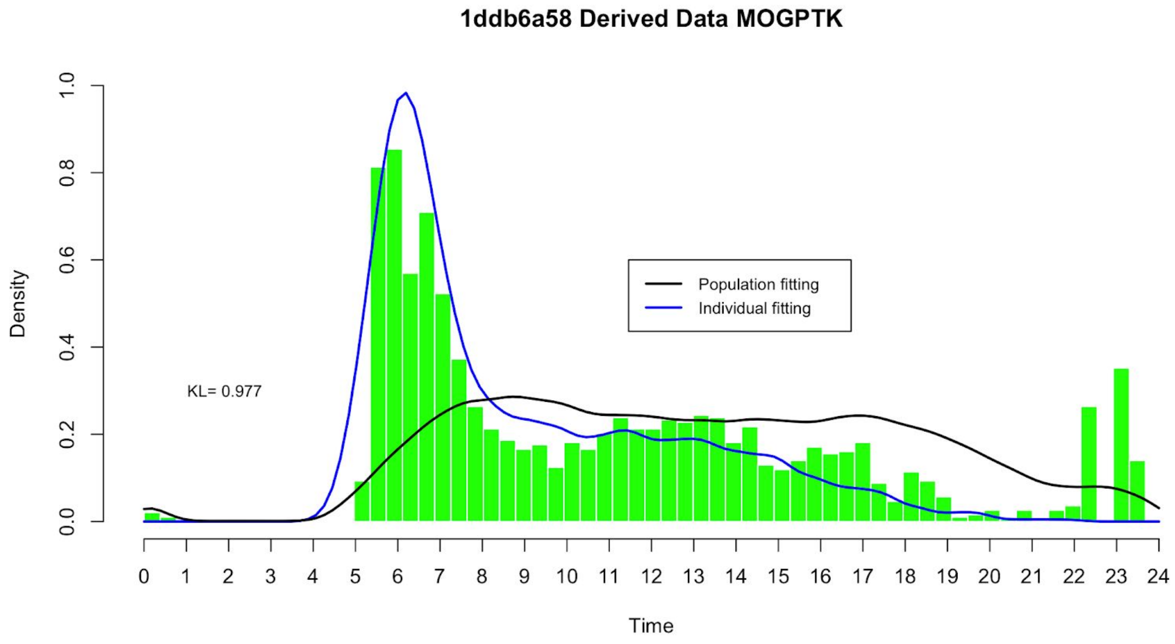

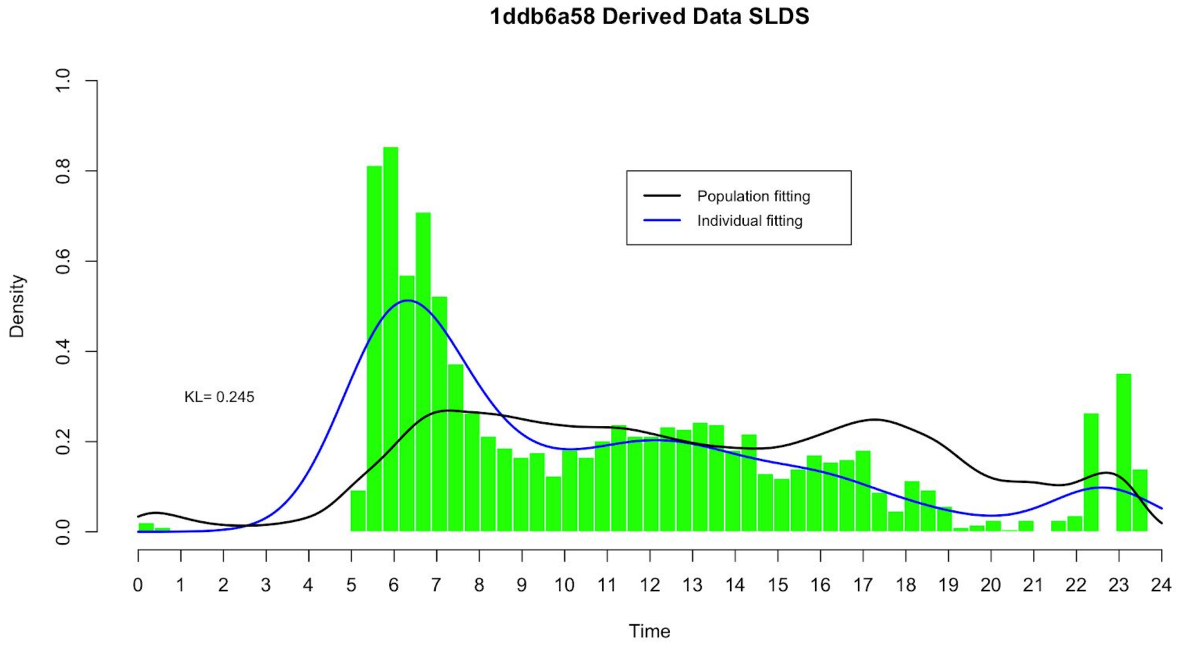
1ddb6a58 Derived Data SLDS
Population fitting
Individual fitting
KL= 0.245
Density
Time
0.0
0.2
0.4
0.6
0.8
1.0
0 1 2 3 4 5 6 7 8 9 10 11 12 13 14 15 16 17 18 19 20 21 22 23 24

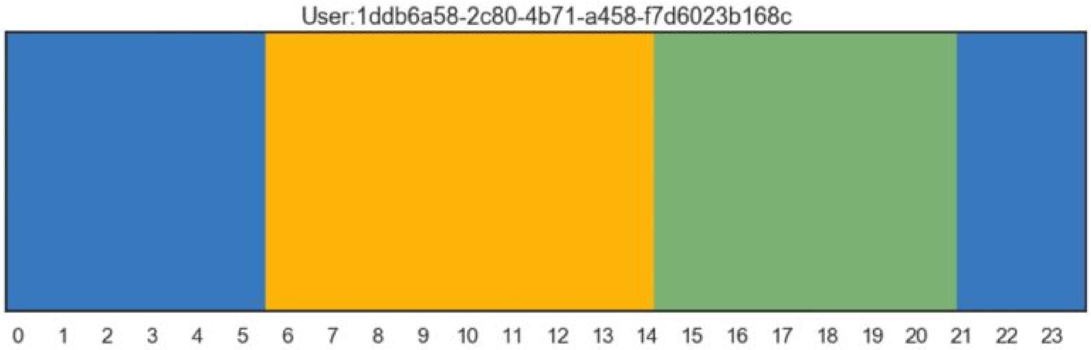
User:1ddb6a58-2c80-4b71-a458-f7d6023b168c
0 1 2 3 4 5 6 7 8 9 10 11 12 13 14 15 16 17 18 19 20 21 22 23

**1ddb6a58 Derived Data MixedVonmises**

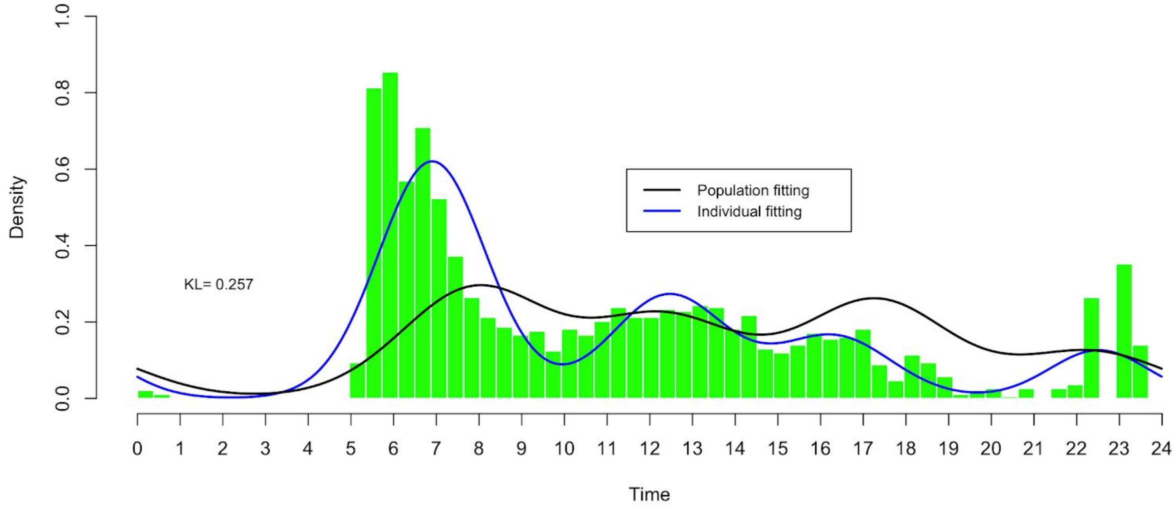


**1ddb6a58 Derived Data HMM**

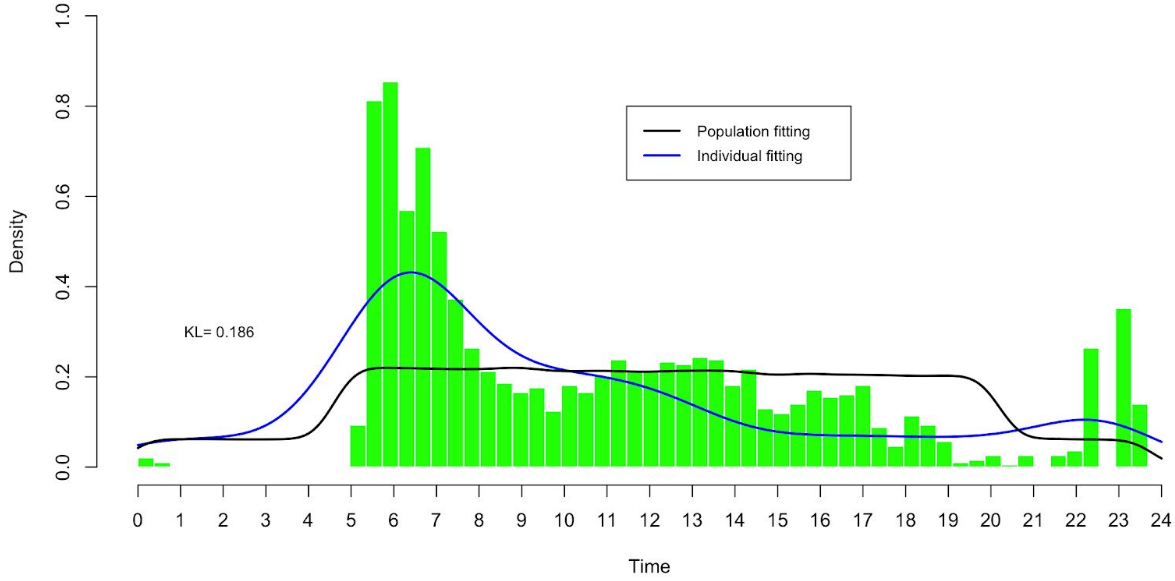

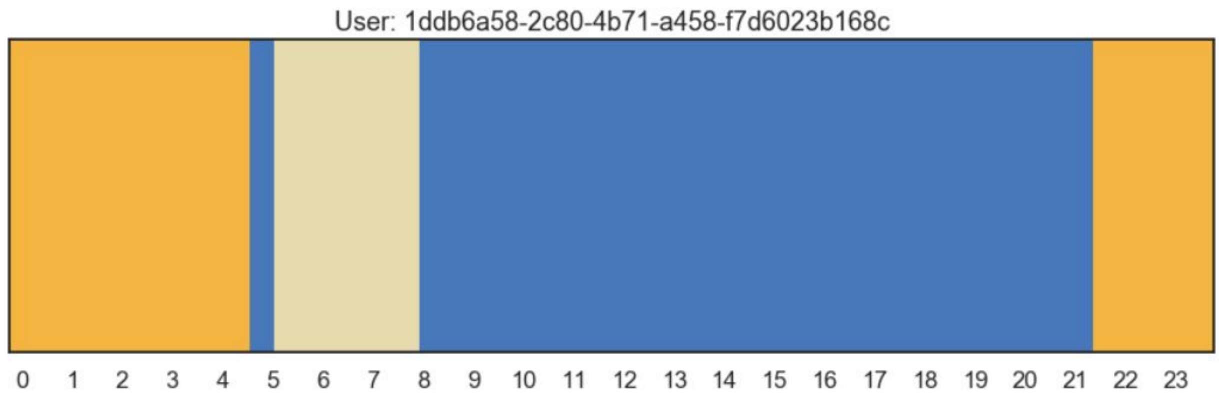


### 42b51b61 Active throughout the day

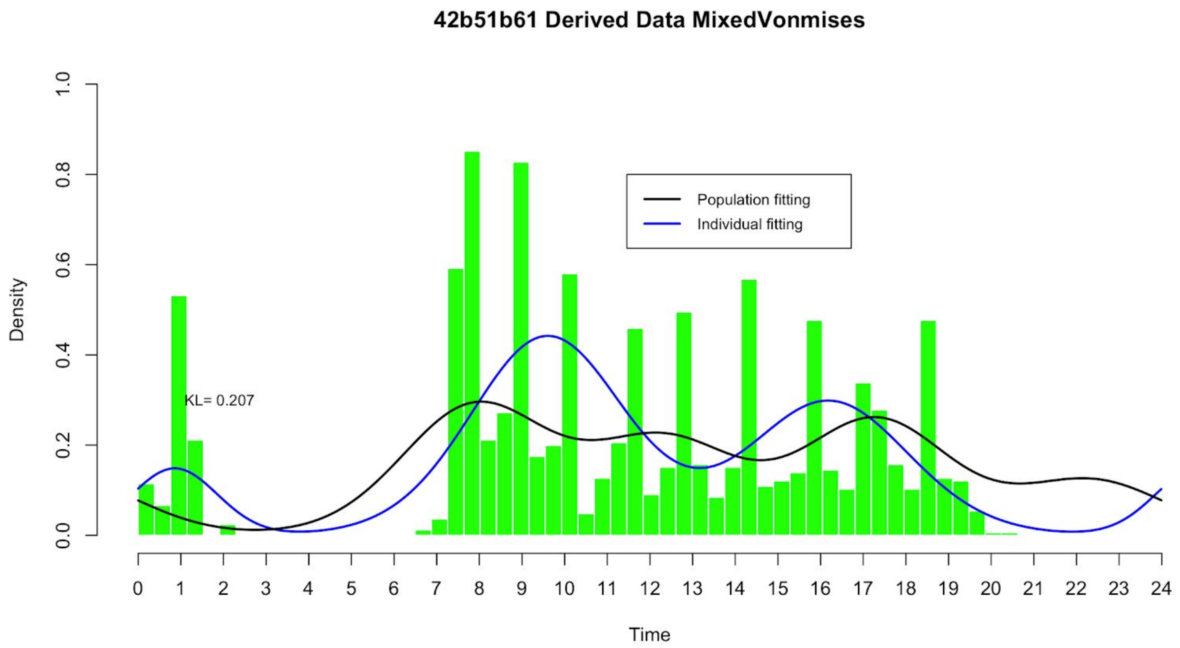

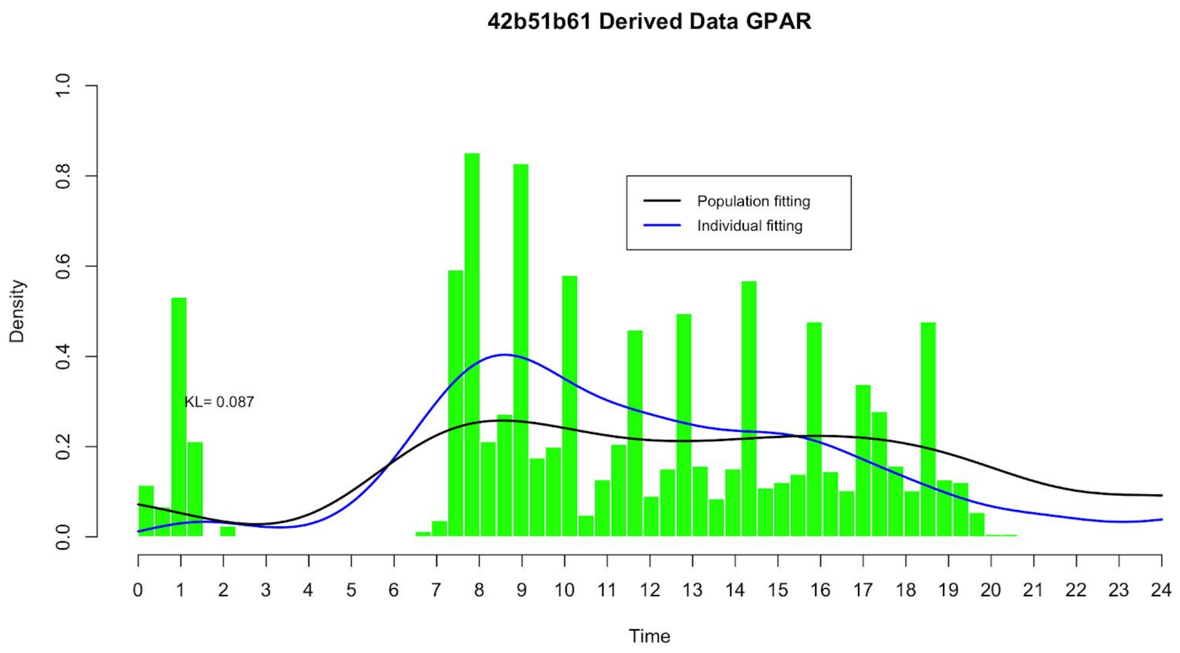
42b51b61 Derived Data GPAR
Population fitting
Individual fitting
KL= 0.087
Density
Time

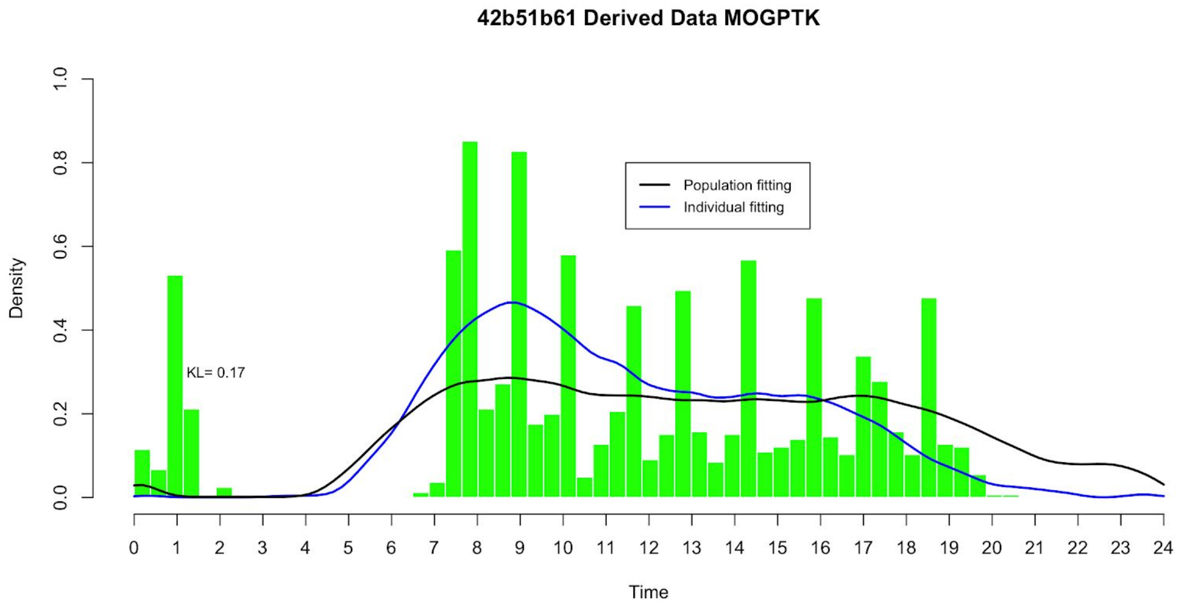
42b51b61 Derived Data MOGPTK
Population fitting
Individual fitting
KL= 0.17
Density
Time

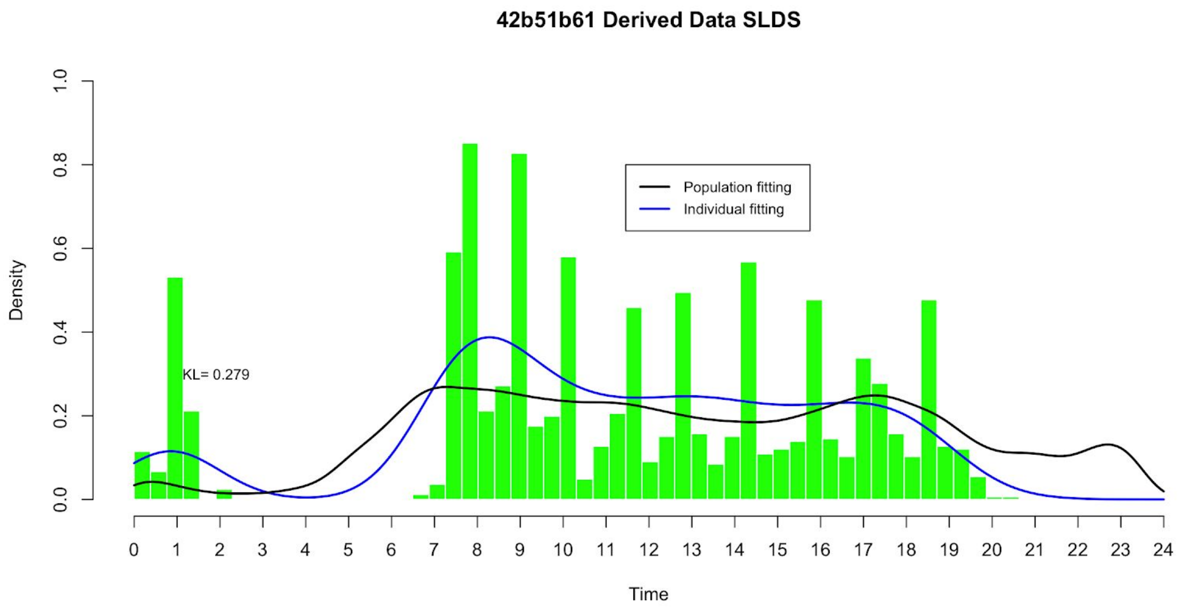
42b51b61 Derived Data SLDS
Population fitting
Individual fitting
KL= 0.279
Density
Time
0.0
0.2
0.4
0.6
0.8
1.0
0 1 2 3 4 5 6 7 8 9 10 11 12 13 14 15 16 17 18 19 20 21 22 23 24

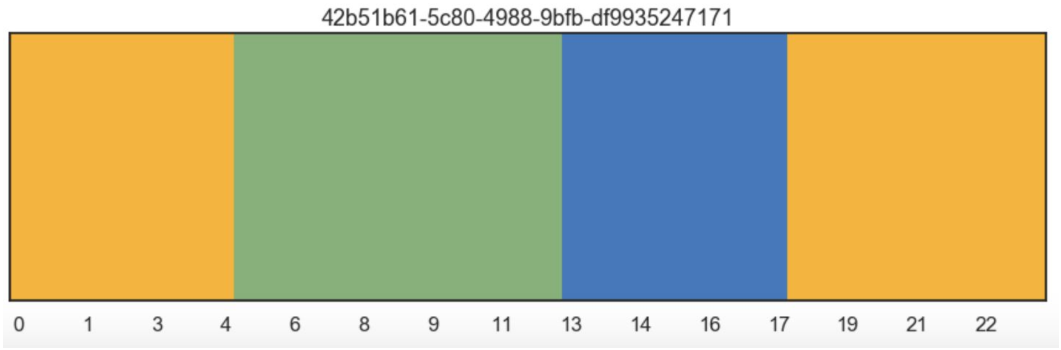
42b51b61-5c80-4988-9bfb-df9935247171
0 1 3 4 6 8 9 11 13 14 16 17 19 21 22

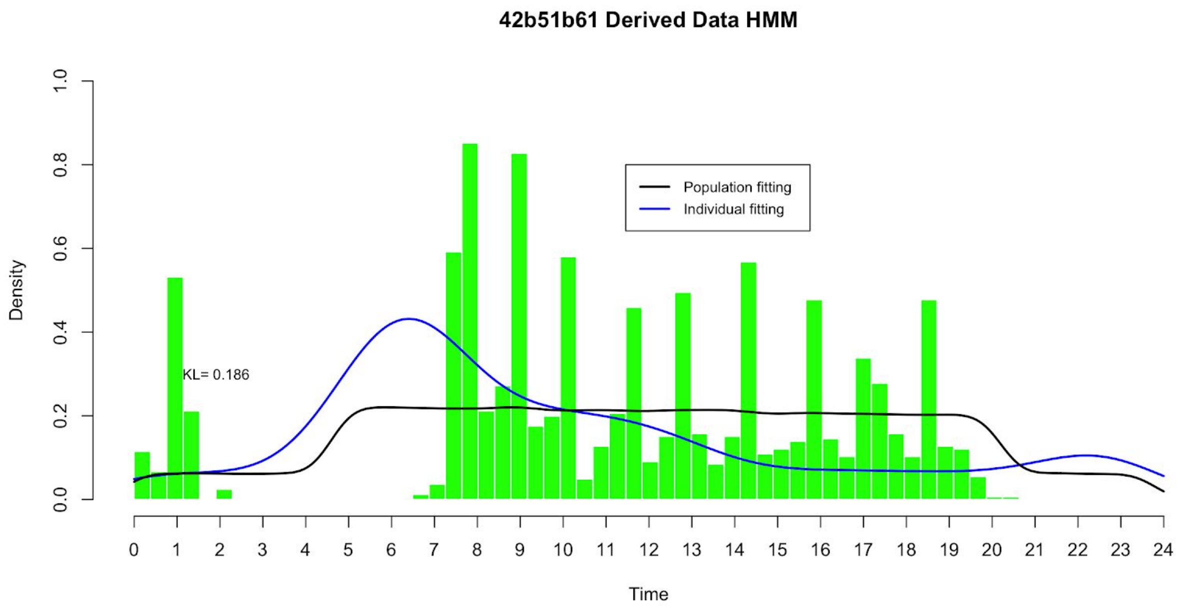
42b51b61 Derived Data HMM
Population fitting
Individual fitting
KL= 0.186
Density
Time
0.0
0.2
0.4
0.6
0.8
1.0
0 1 2 3 4 5 6 7 8 9 10 11 12 13 14 15 16 17 18 19 20 21 22 23 24

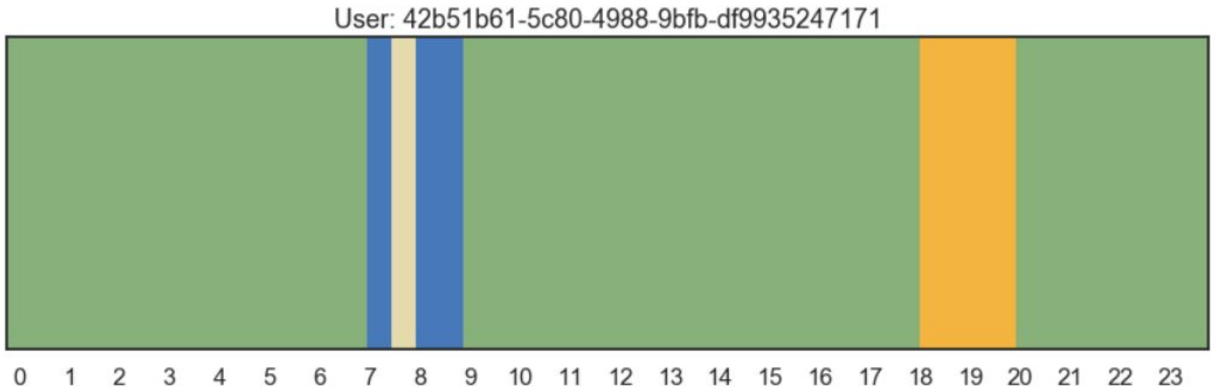
User: 42b51b61-5c80-4988-9bfb-df9935247171
0 1 2 3 4 5 6 7 8 9 10 11 12 13 14 15 16 17 18 19 20 21 22 23

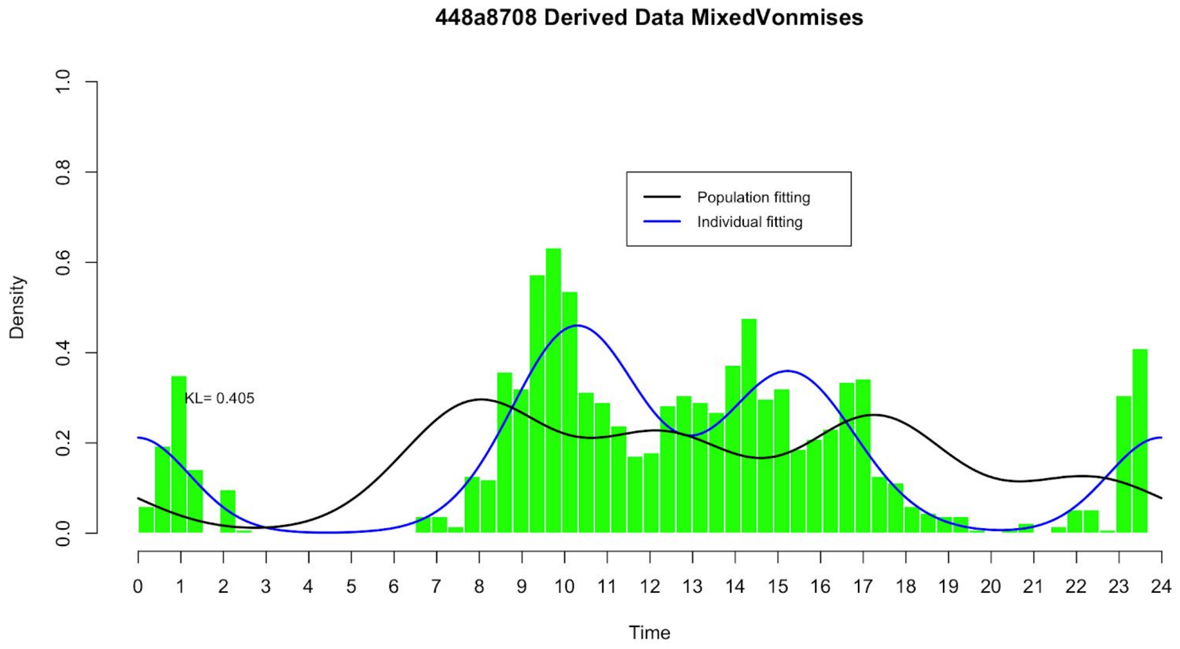

448a8708 Derived Data MixedVonmises
Population fitting
Individual fitting
KL= 0.405
Density
Time
0.0
0.2
0.4
0.6
0.8
1.0
0 1 2 3 4 5 6 7 8 9 10 11 12 13 14 15 16 17 18 19 20 21 22 23 24

### 9821cc46 Active later than most

**9821cc46 Derived Data MixedVonmises**

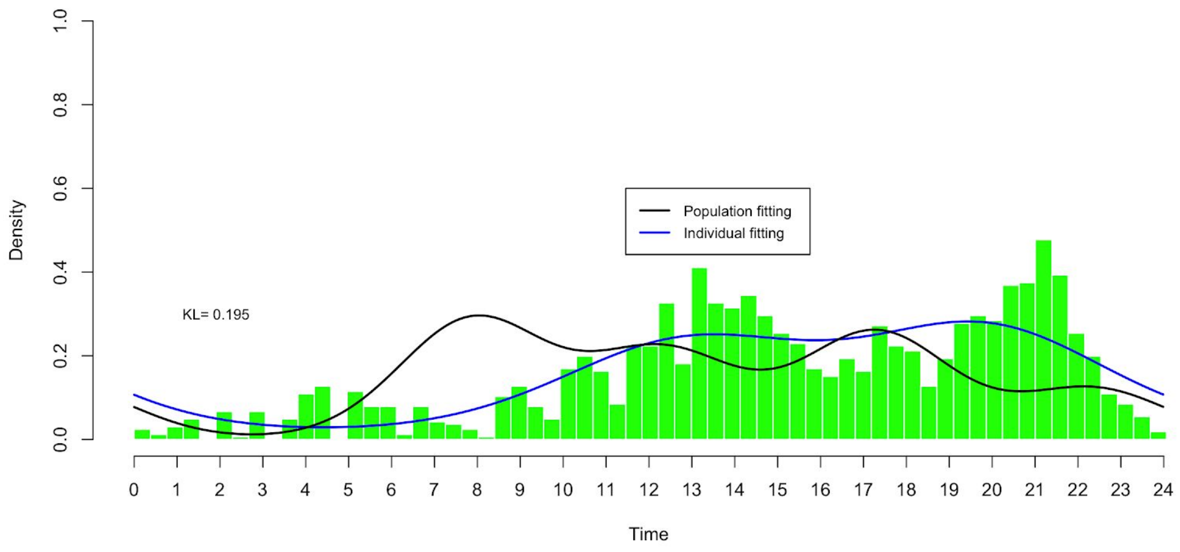


**9821cc46 Derived Data GPAR**

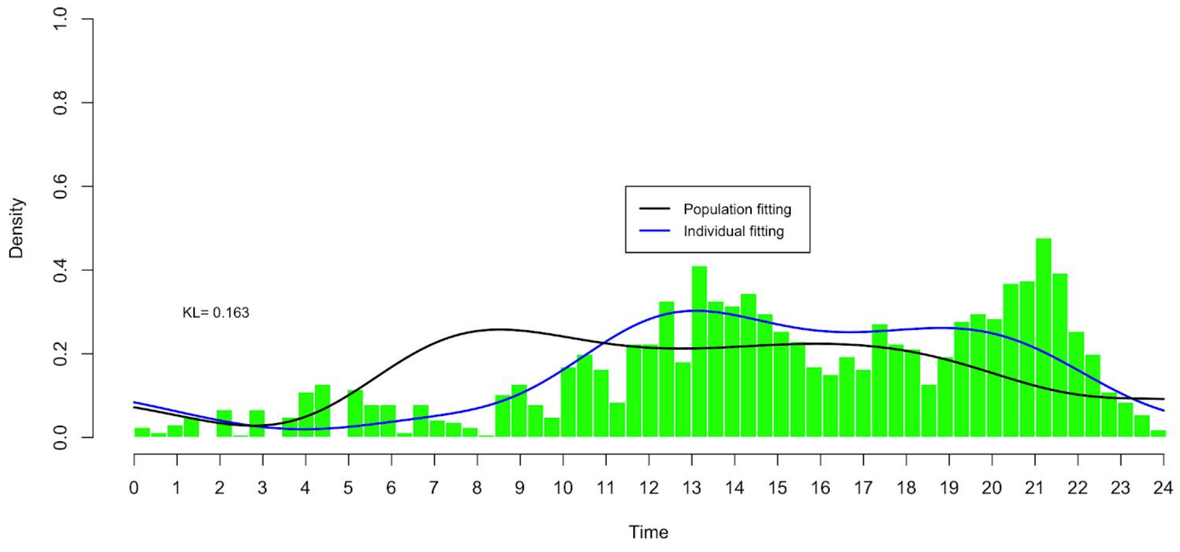

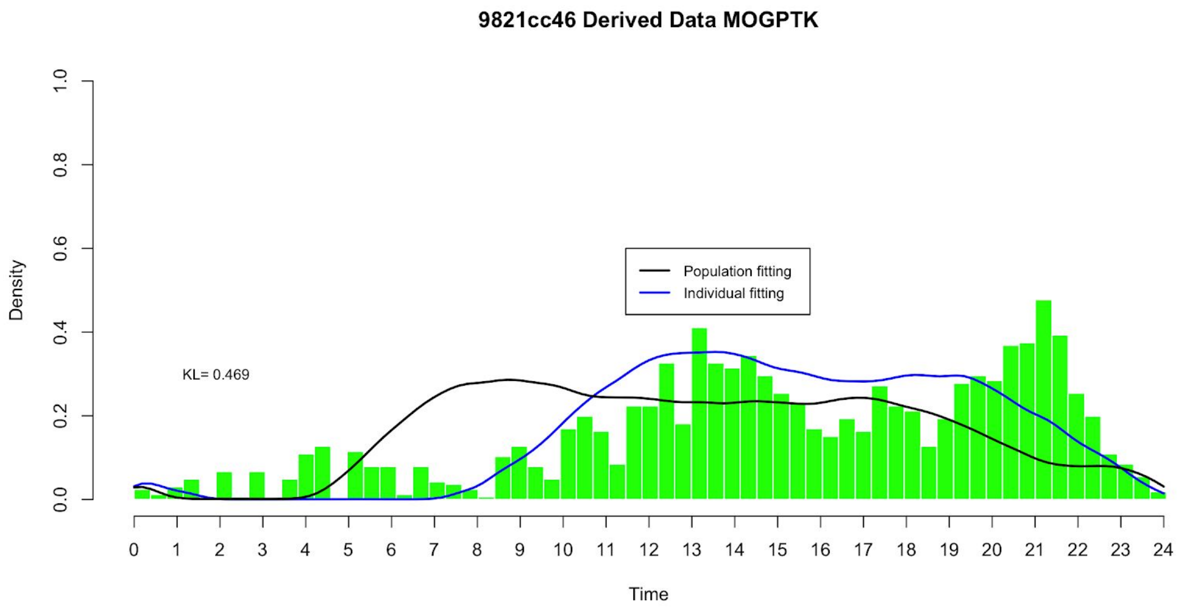
9821cc46 Derived Data MOGPTK
Density
Time
Population fitting
Individual fitting
KL= 0.469

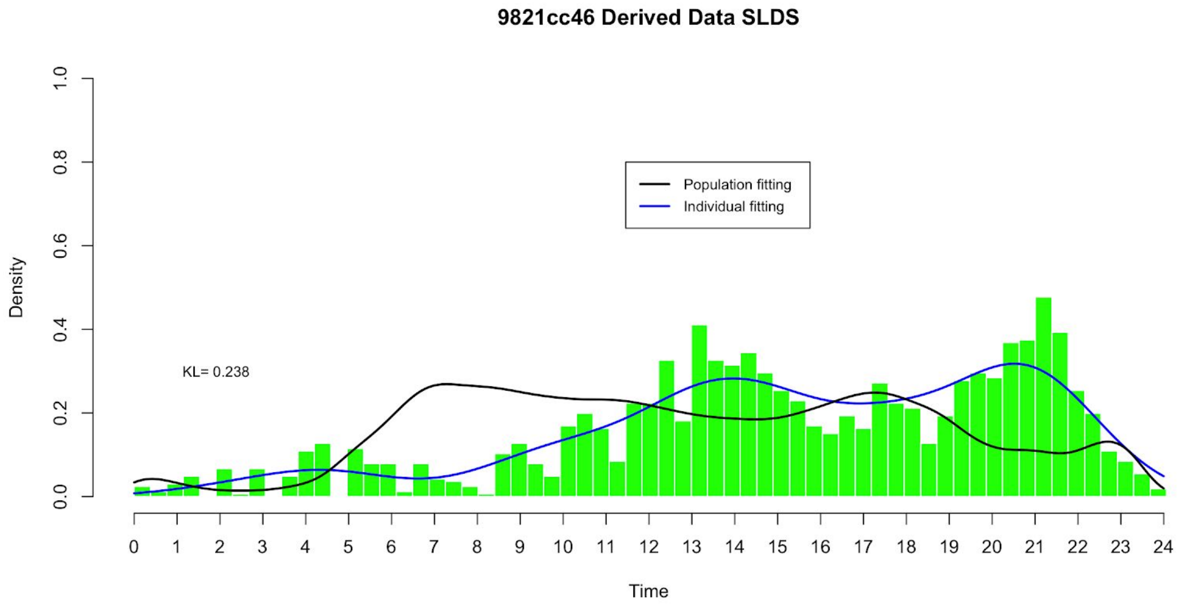
9821cc46 Derived Data SLDS
Density
Time
Population fitting
Individual fitting
KL= 0.238

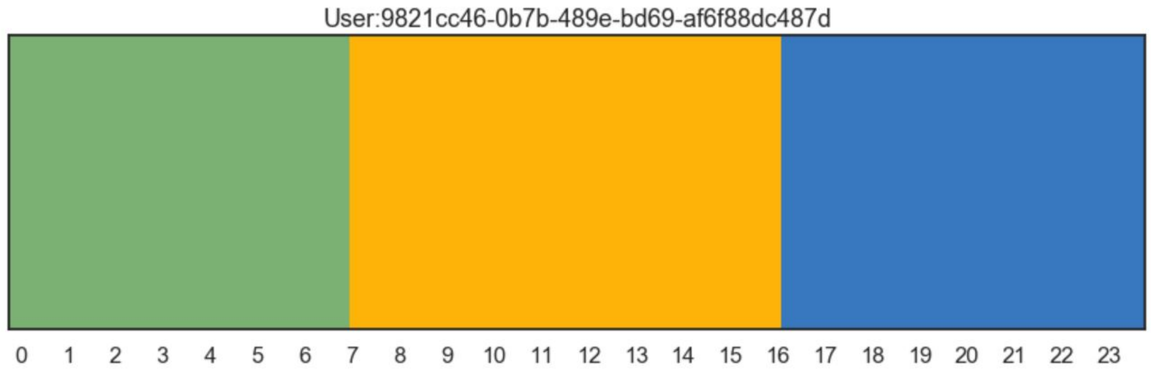


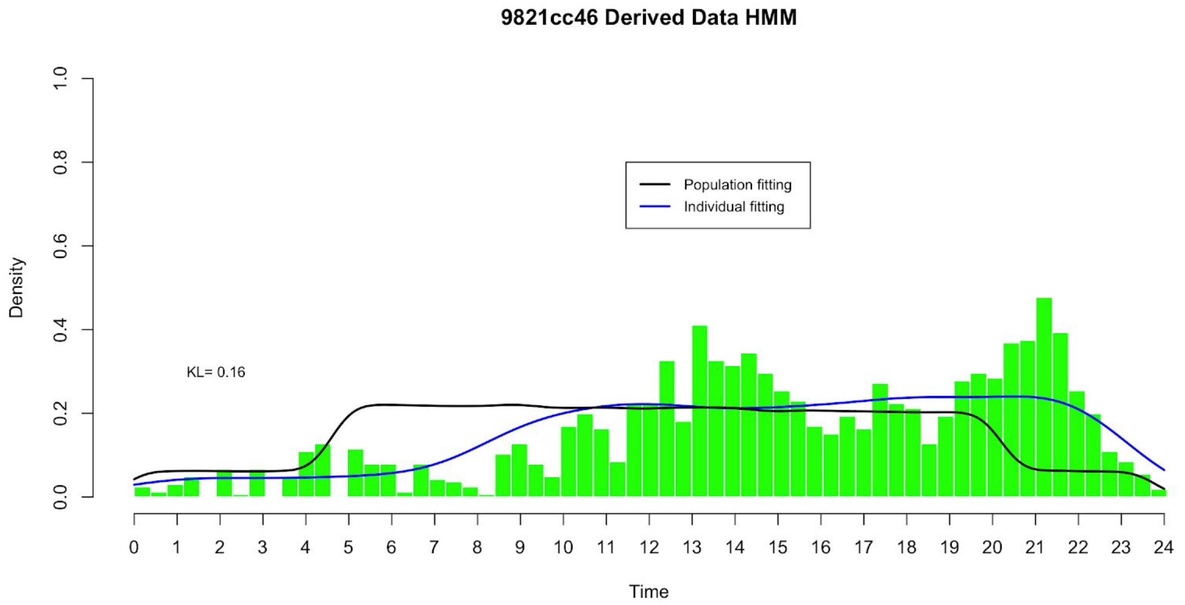


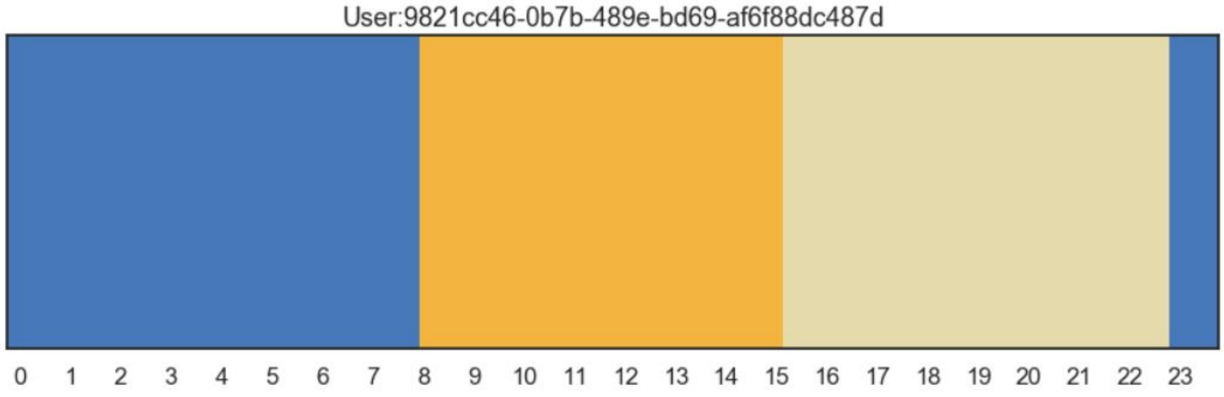


2b2cbc5d Active more in the early part of the day

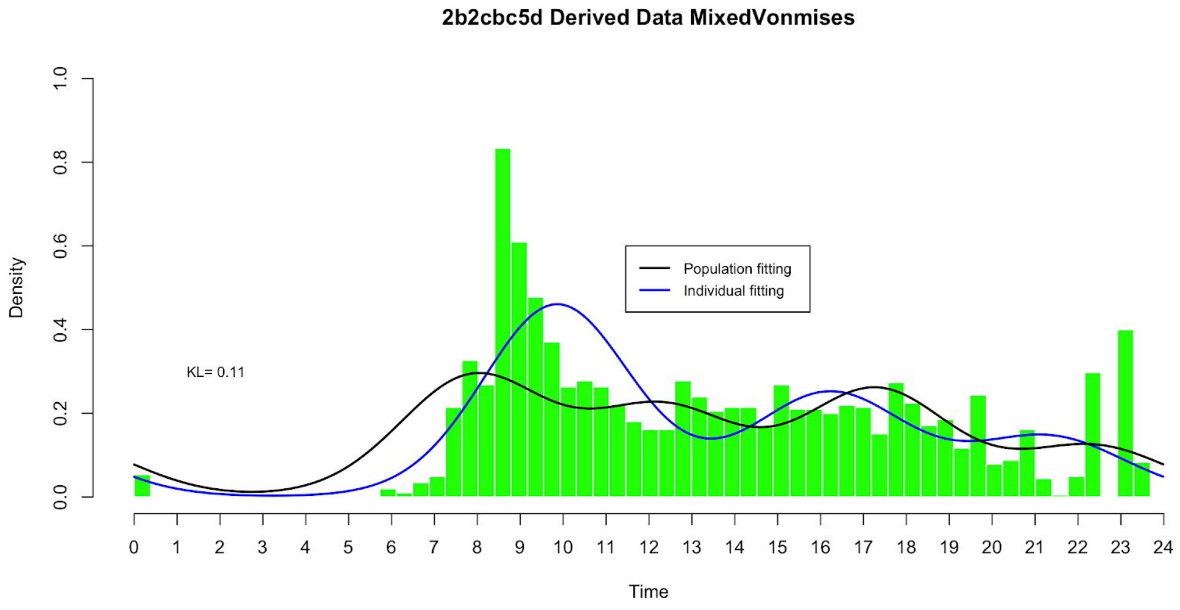
2b2cbc5d Derived Data MixedVonmises
Population fitting
Individual fitting
KL= 0.11
Density
Time
0.0
0.2
0.4
0.6
0.8
1.0
0 1 2 3 4 5 6 7 8 9 10 11 12 13 14 15 16 17 18 19 20 21 22 23 24

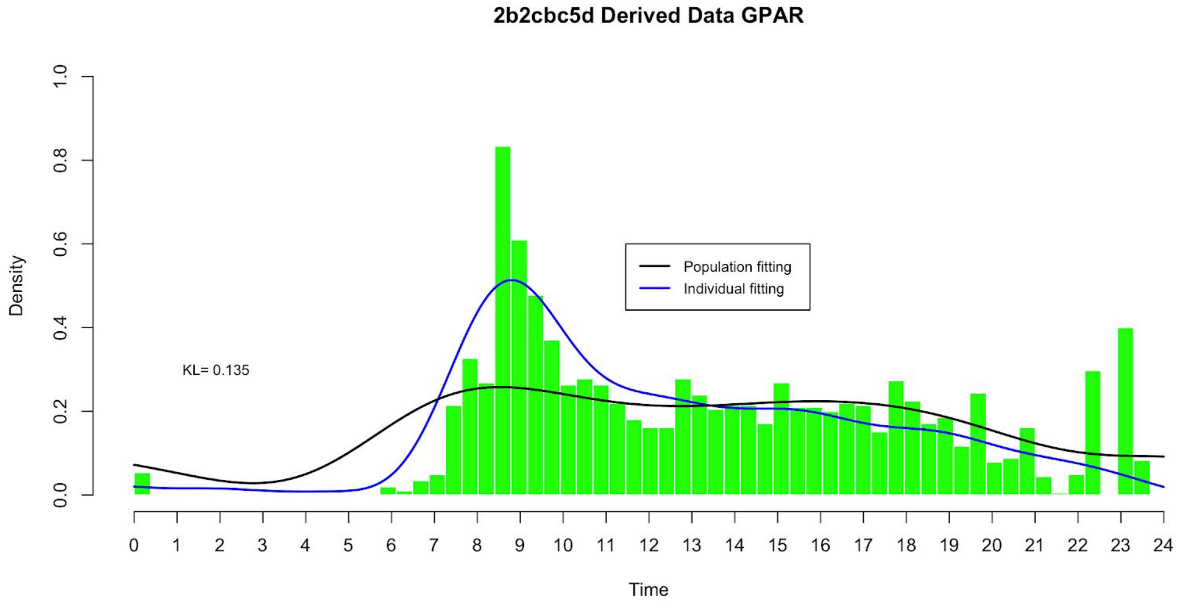
2b2cbc5d Derived Data GPAR
Population fitting
Individual fitting
KL= 0.135
Density
Time
0.0
0.2
0.4
0.6
0.8
1.0
0 1 2 3 4 5 6 7 8 9 10 11 12 13 14 15 16 17 18 19 20 21 22 23 24

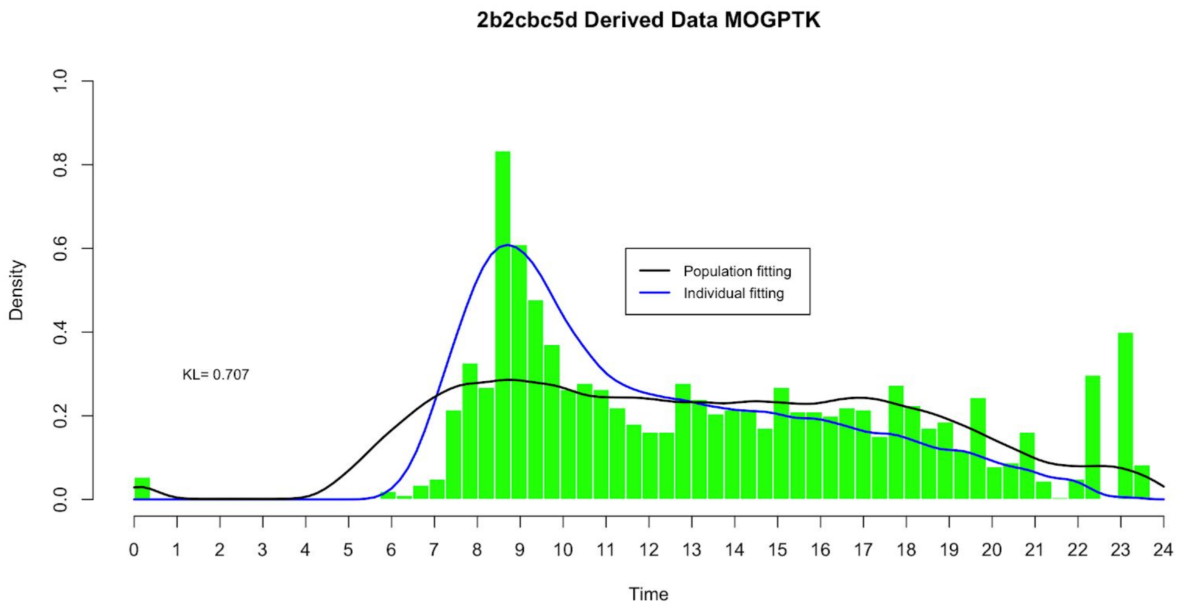
2b2cbc5d Derived Data MOGPTK
Population fitting
Individual fitting
KL= 0.707
Density
Time
0.0
0.2
0.4
0.6
0.8
1.0
0 1 2 3 4 5 6 7 8 9 10 11 12 13 14 15 16 17 18 19 20 21 22 23 24

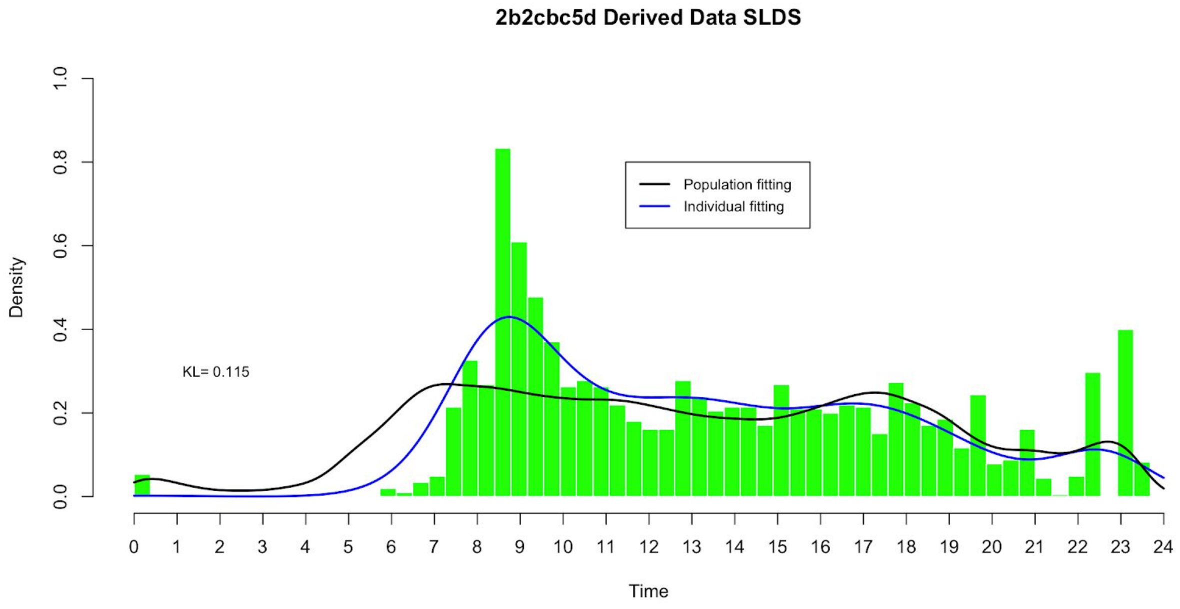
2b2cbc5d Derived Data SLDS
Population fitting
Individual fitting
KL= 0.115
Density
Time
0.0
0.2
0.4
0.6
0.8
1.0
0 1 2 3 4 5 6 7 8 9 10 11 12 13 14 15 16 17 18 19 20 21 22 23 24

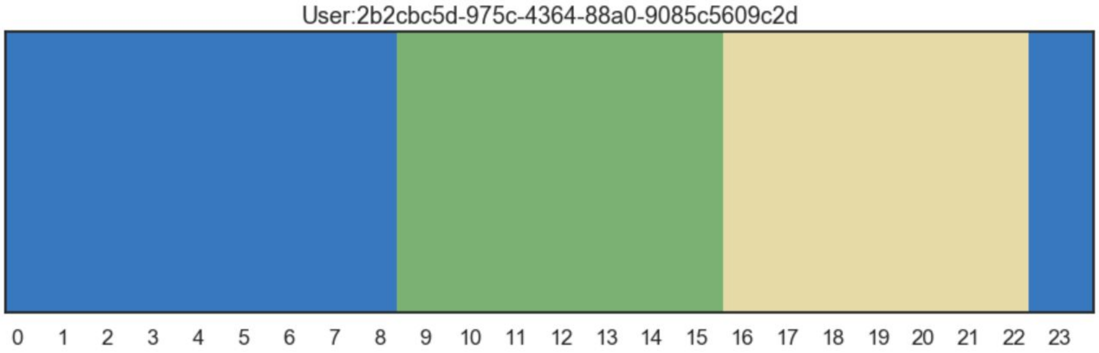
User:2b2cbc5d-975c-4364-88a0-9085c5609c2d
0 1 2 3 4 5 6 7 8 9 10 11 12 13 14 15 16 17 18 19 20 21 22 23

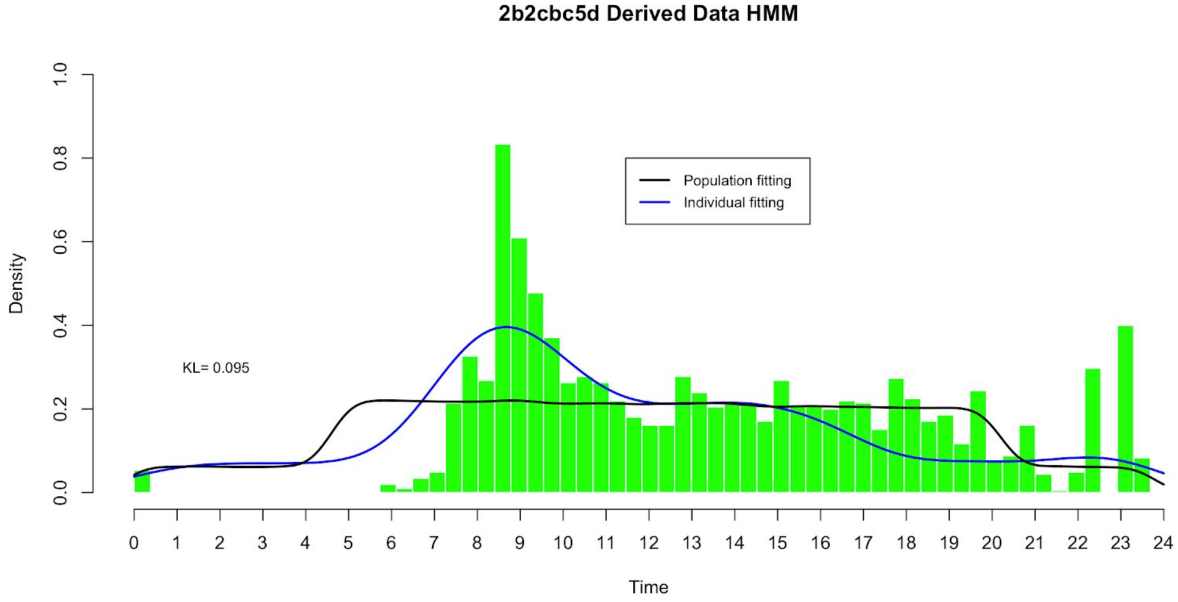
2b2cbc5d Derived Data HMM
Population fitting
Individual fitting
KL= 0.095
Density
0.0 0.2 0.4 0.6 0.8 1.0
0 1 2 3 4 5 6 7 8 9 10 11 12 13 14 15 16 17 18 19 20 21 22 23 24
Time

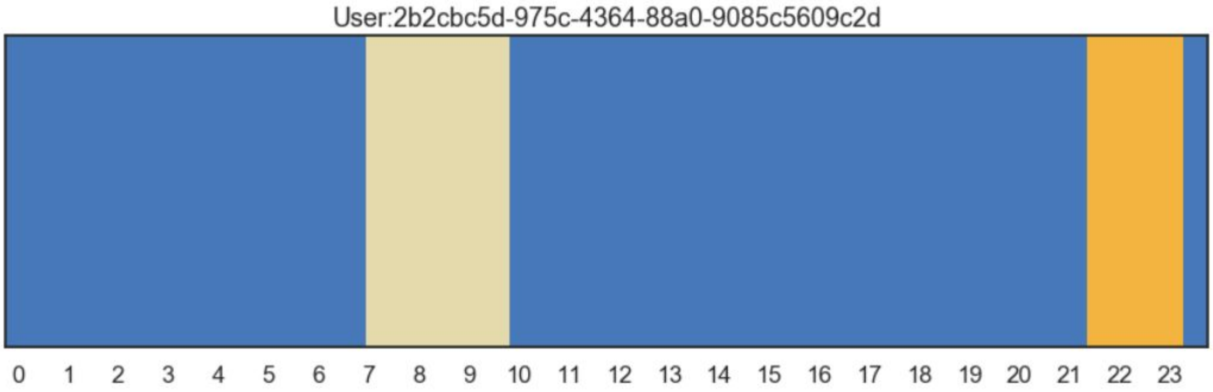
User:2b2cbc5d-975c-4364-88a0-9085c5609c2d
0 1 2 3 4 5 6 7 8 9 10 11 12 13 14 15 16 17 18 19 20 21 22 23

### 7d439102 Quieter in the middle part of the day

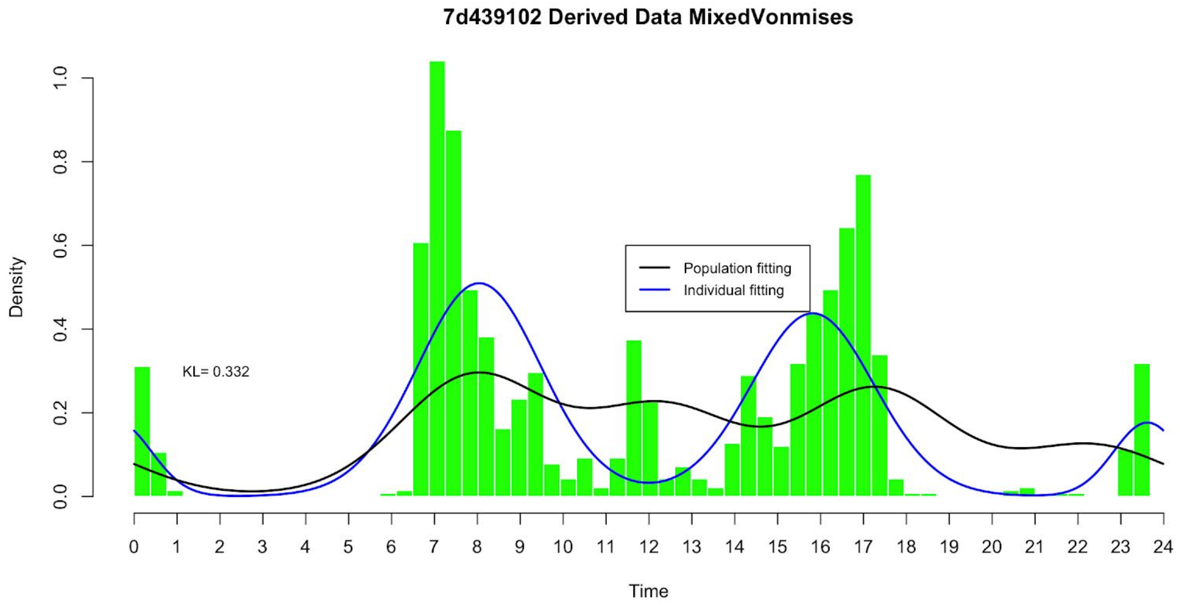


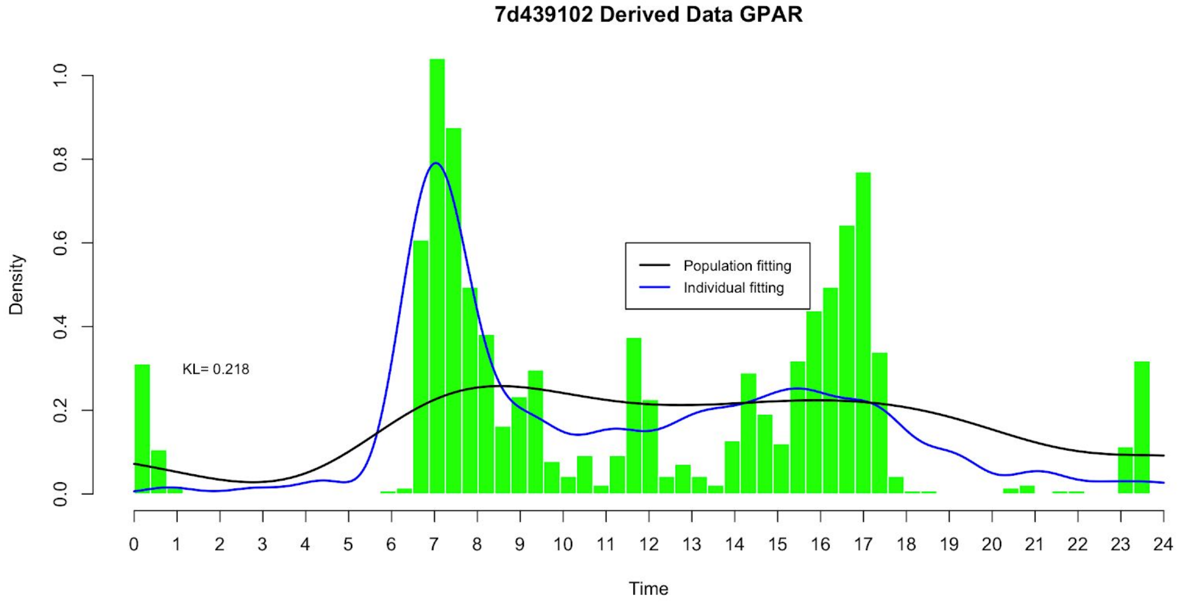

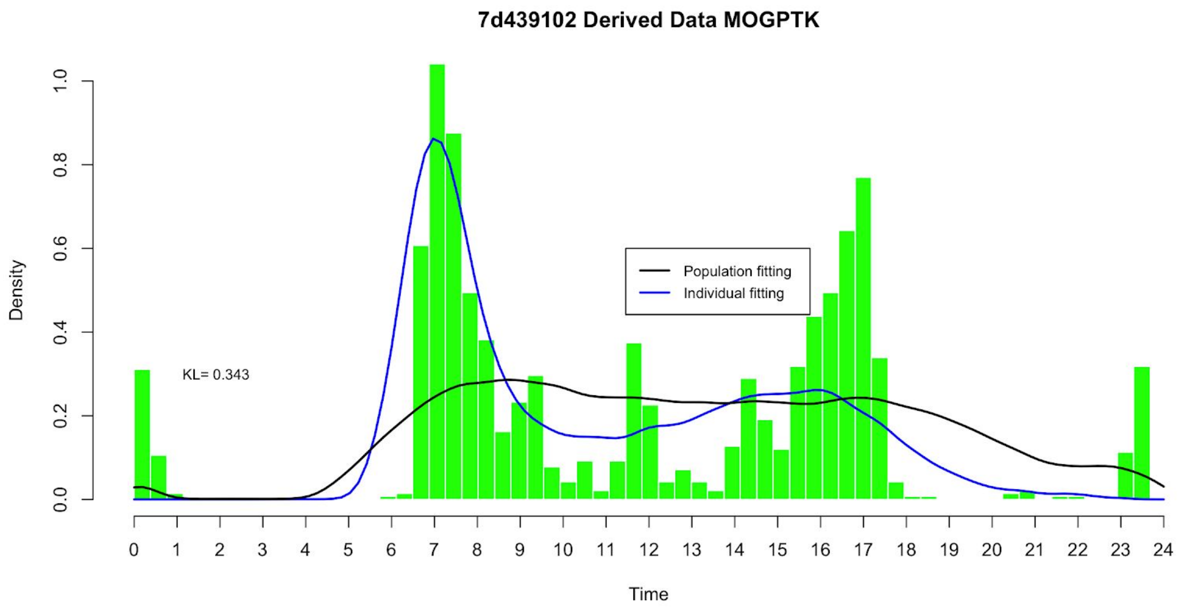

7d439102 Derived Data MOGPTK
Population fitting
Individual fitting
KL= 0.343
Density
Time


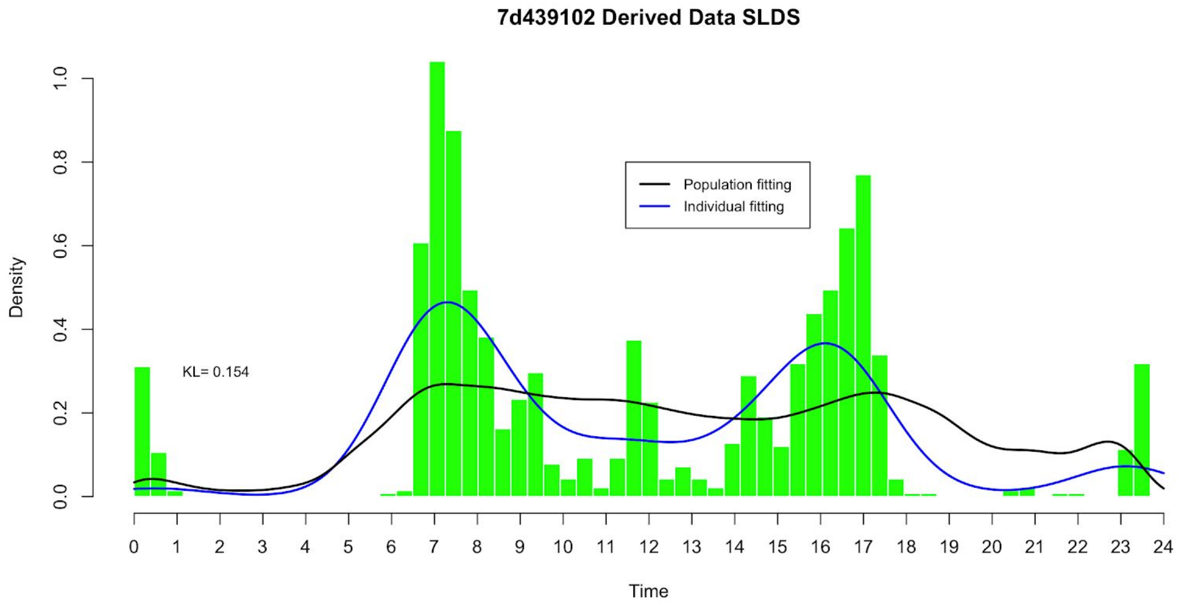

7d439102 Derived Data SLDS
Population fitting
Individual fitting
KL= 0.154
Density
Time


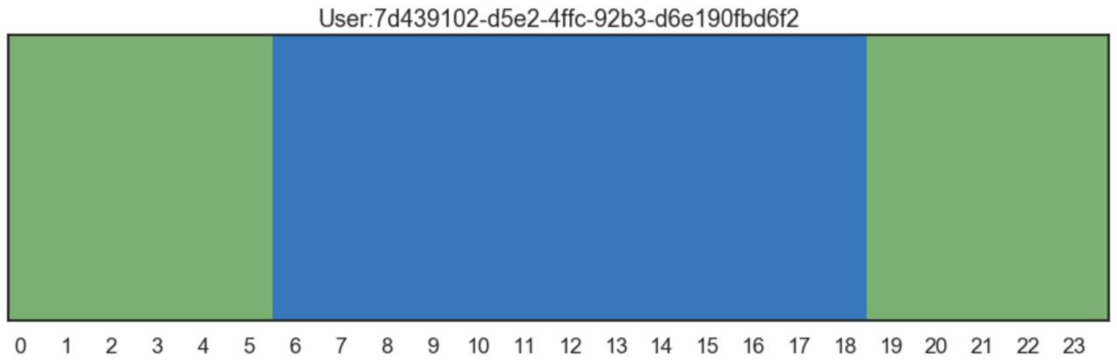

User:7d439102-d5e2-4ffc-92b3-d6e190fbd6f2

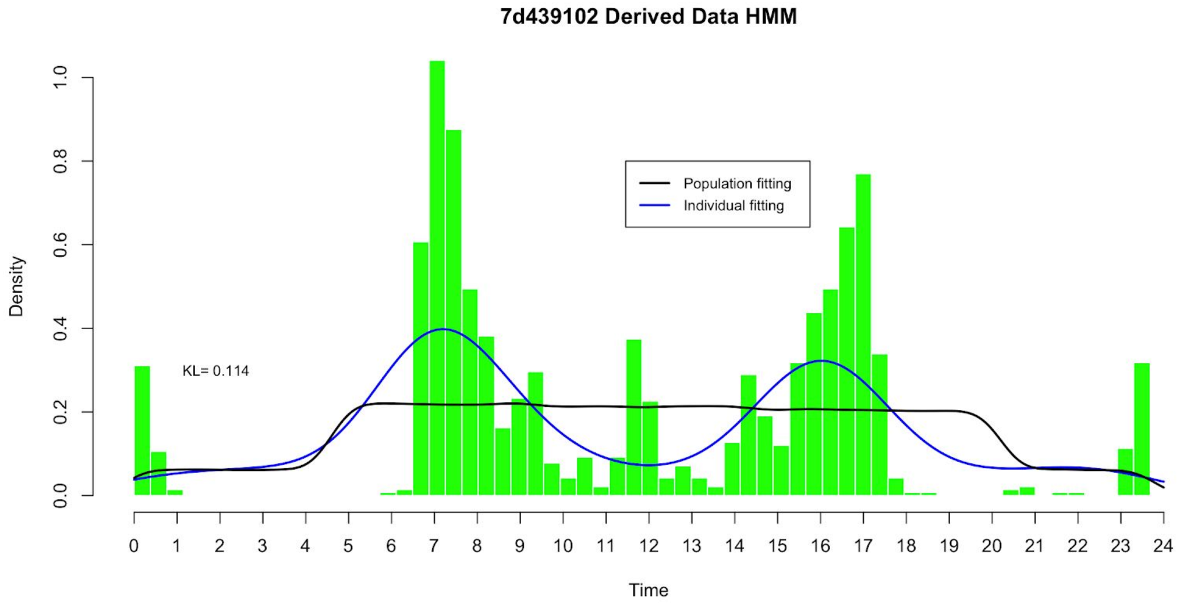
7d439102 Derived Data HMM
Density
Time
Population fitting
Individual fitting
KL= 0.114
0.0
0.2
0.4
0.6
0.8
1.0
0 1 2 3 4 5 6 7 8 9 10 11 12 13 14 15 16 17 18 19 20 21 22 23 24

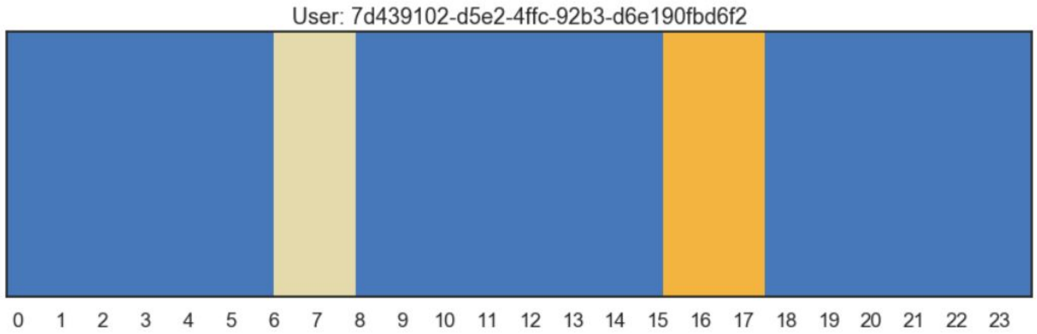
User: 7d439102-d5e2-4ffc-92b3-d6e190fbd6f2
0 1 2 3 4 5 6 7 8 9 10 11 12 13 14 15 16 17 18 19 20 21 22 23

### Individual fitting on primary data

#### 9821cc46 Active later than most

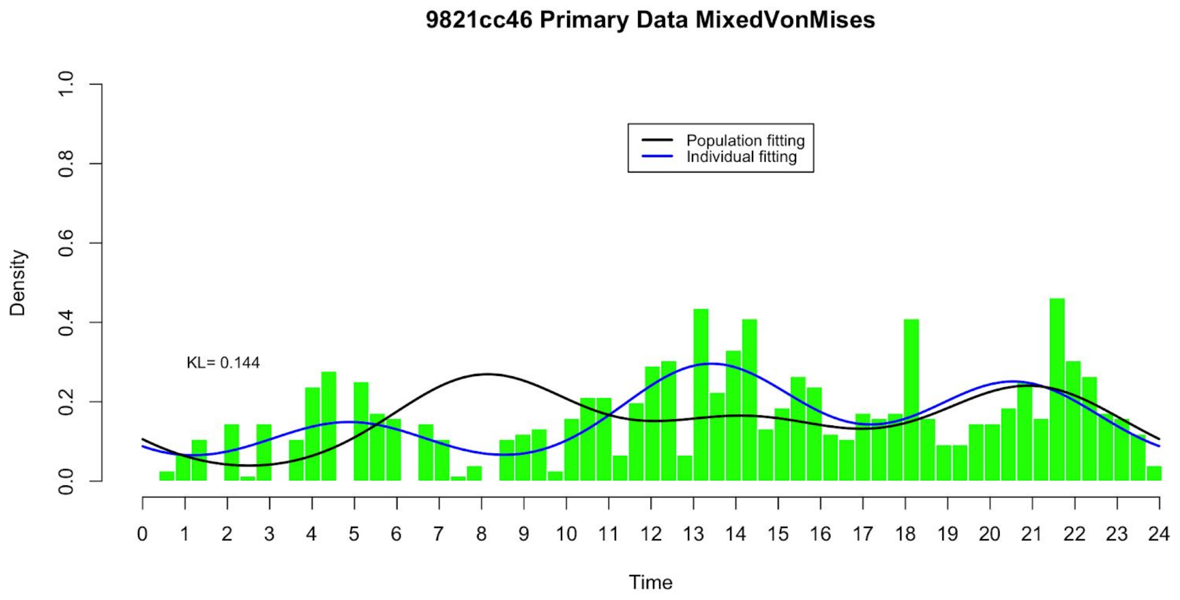


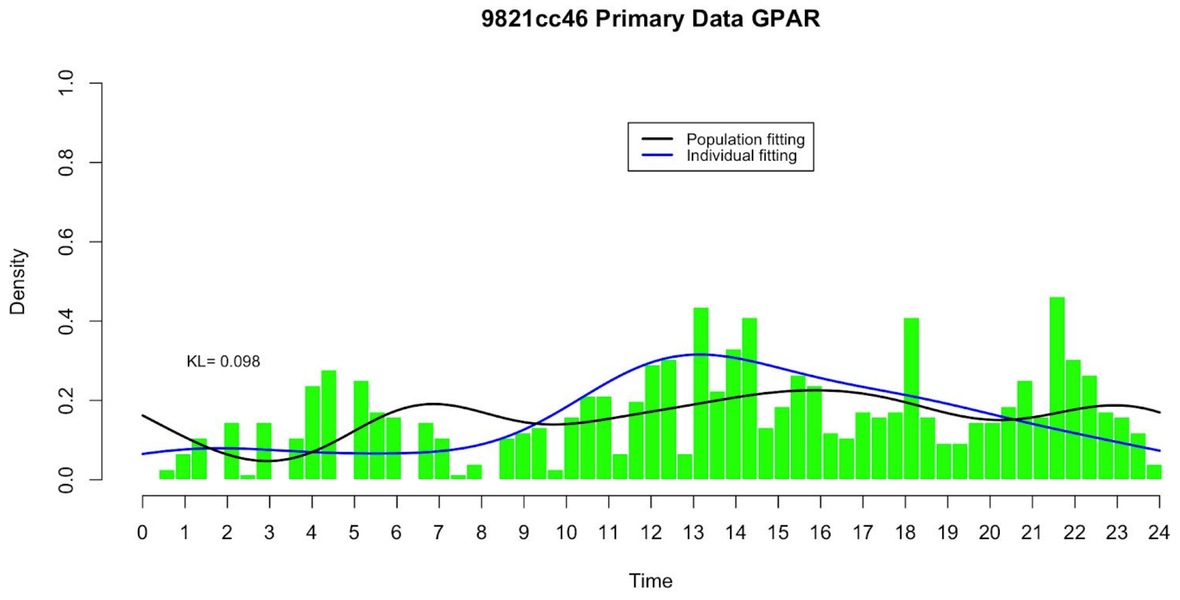

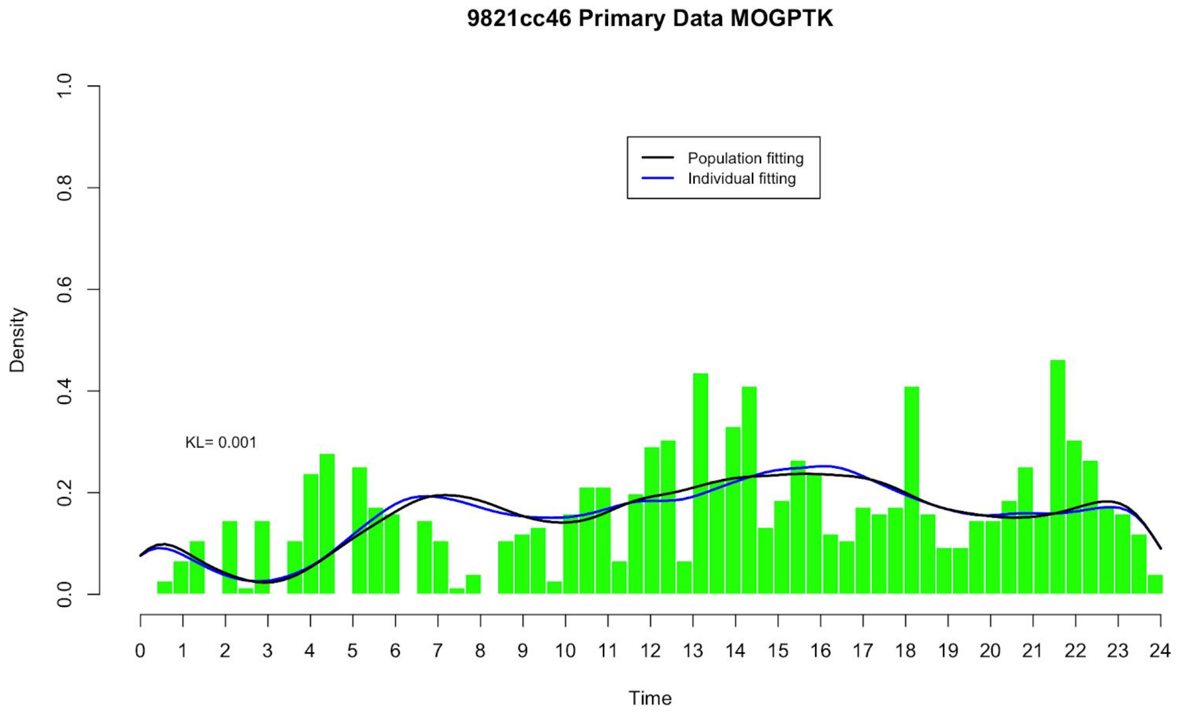

9821cc46 Primary Data MOGPTK
Density
Time
Population fitting
Individual fitting
KL= 0.001


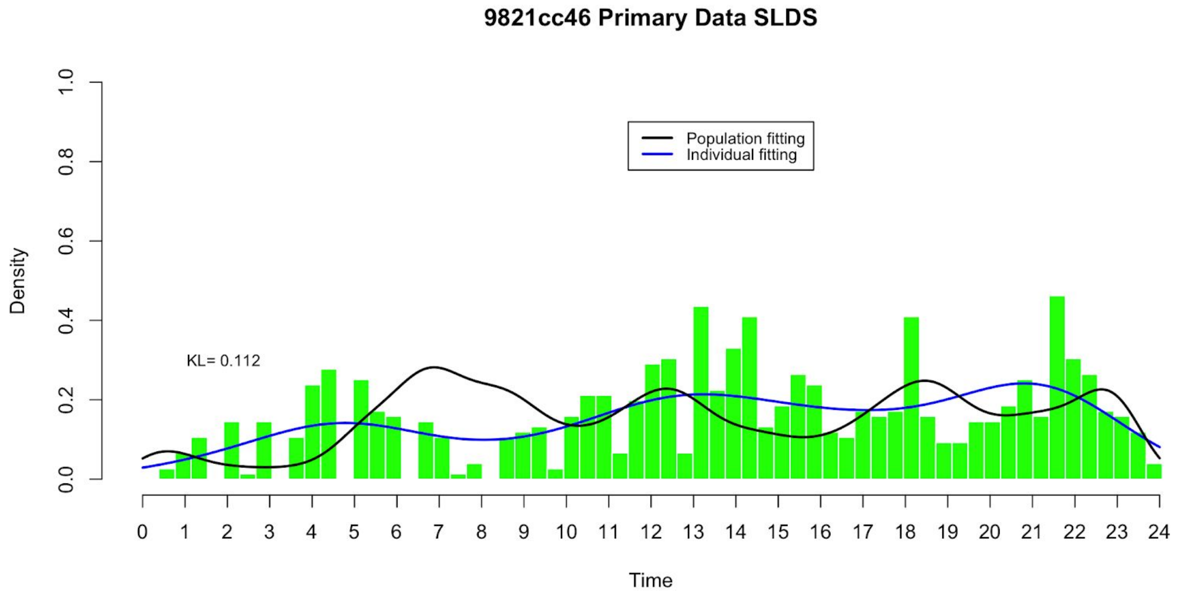

9821cc46 Primary Data SLDS
Density
Time
Population fitting
Individual fitting
KL= 0.112

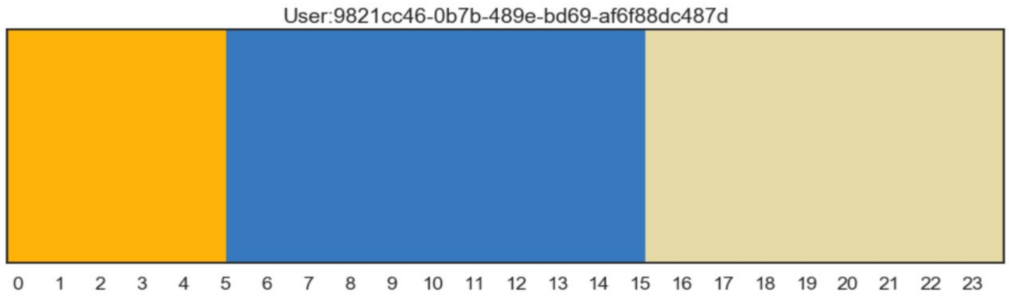
User:9821cc46-0b7b-489e-bd69-af6f88dc487d
0 1 2 3 4 5 6 7 8 9 10 11 12 13 14 15 16 17 18 19 20 21 22 23

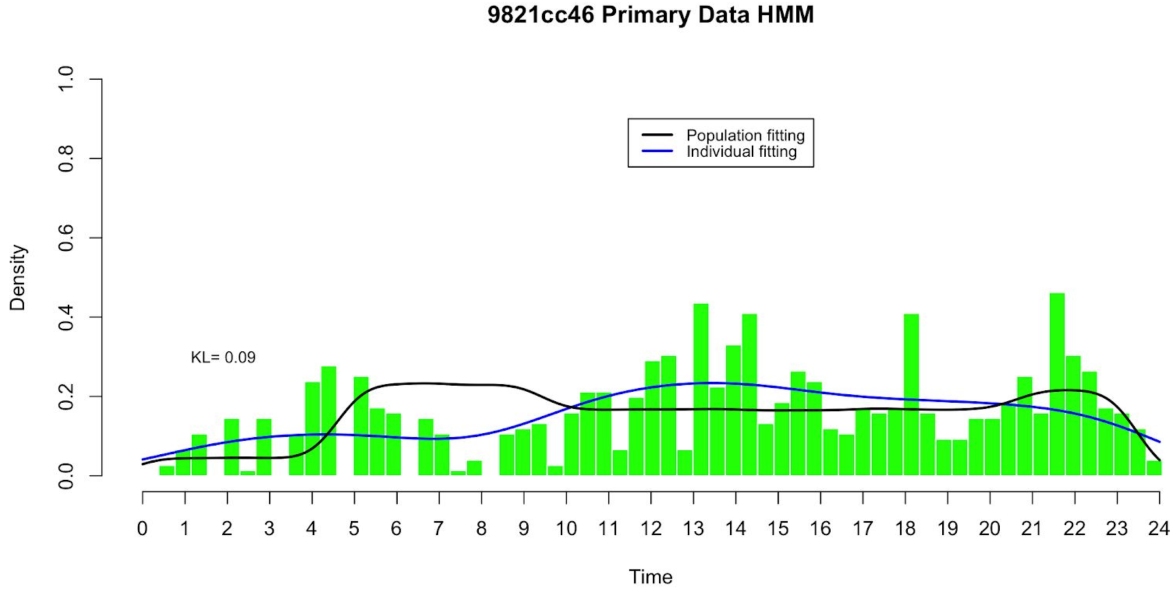
9821cc46 Primary Data HMM
Population fitting
Individual fitting
KL= 0.09
Density
0.0 0.2 0.4 0.6 0.8 1.0
0 1 2 3 4 5 6 7 8 9 10 11 12 13 14 15 16 17 18 19 20 21 22 23 24
Time

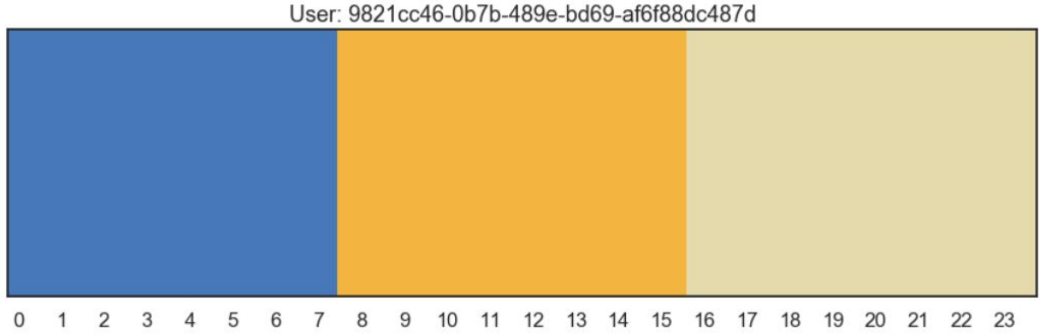
User: 9821cc46-0b7b-489e-bd69-af6f88dc487d
0 1 2 3 4 5 6 7 8 9 10 11 12 13 14 15 16 17 18 19 20 21 22 23

### 2b2cbc5d Active more in the early part of the day

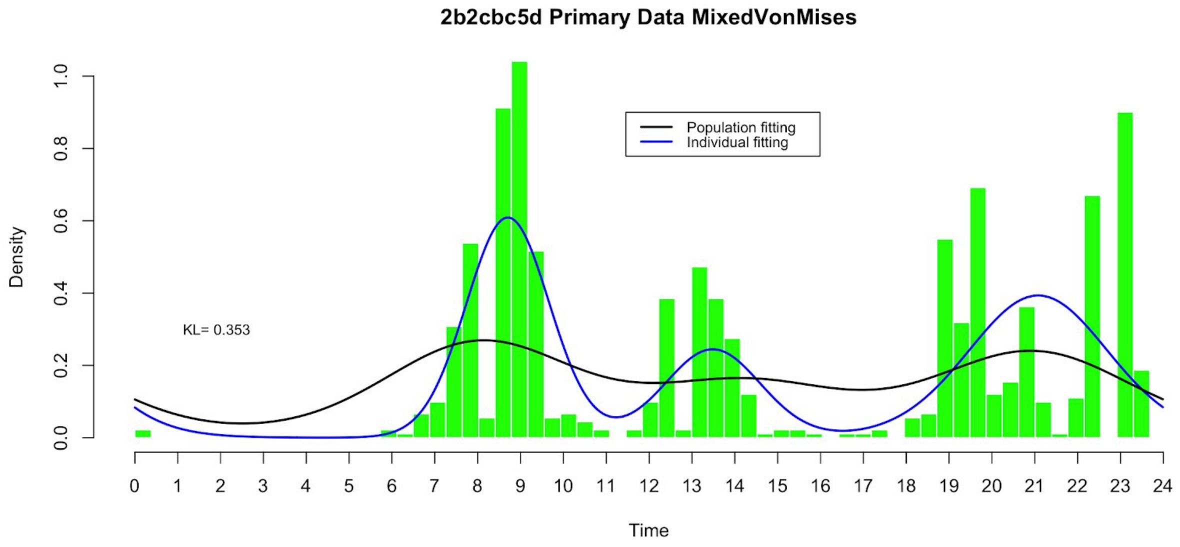


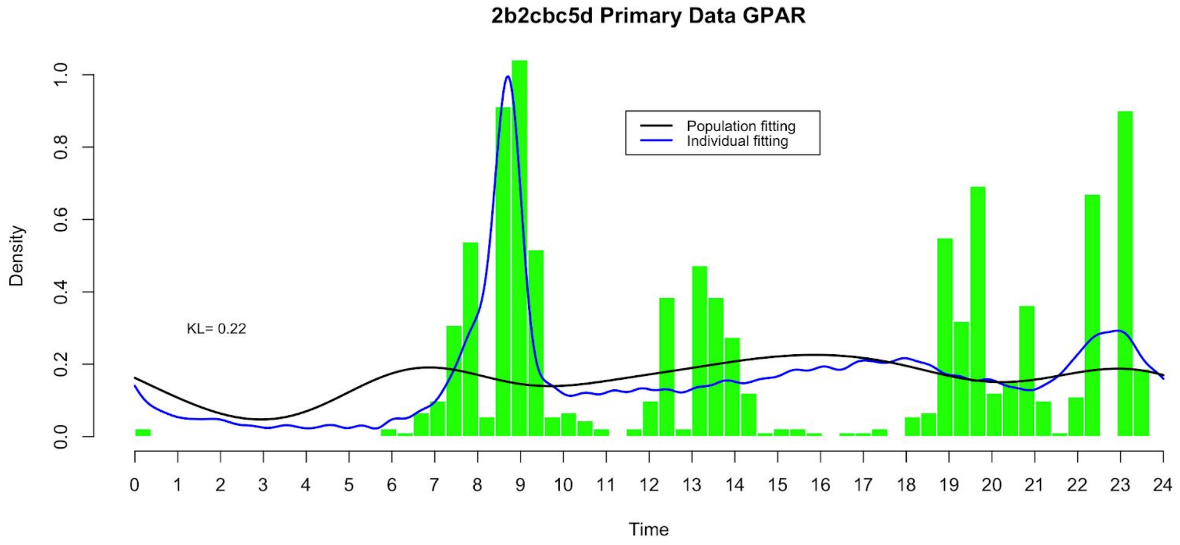

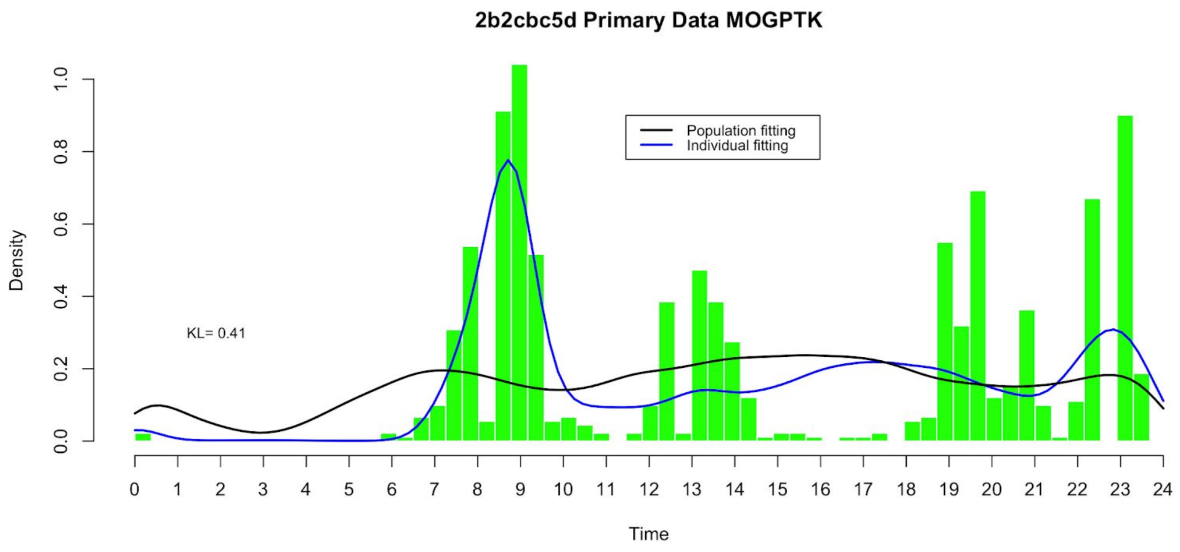
2b2cbc5d Primary Data MOGPTK
Population fitting
Individual fitting
KL= 0.41
Density
0.0
0.2
0.4
0.6
0.8
1.0
0 1 2 3 4 5 6 7 8 9 10 11 12 13 14 15 16 17 18 19 20 21 22 23 24
Time

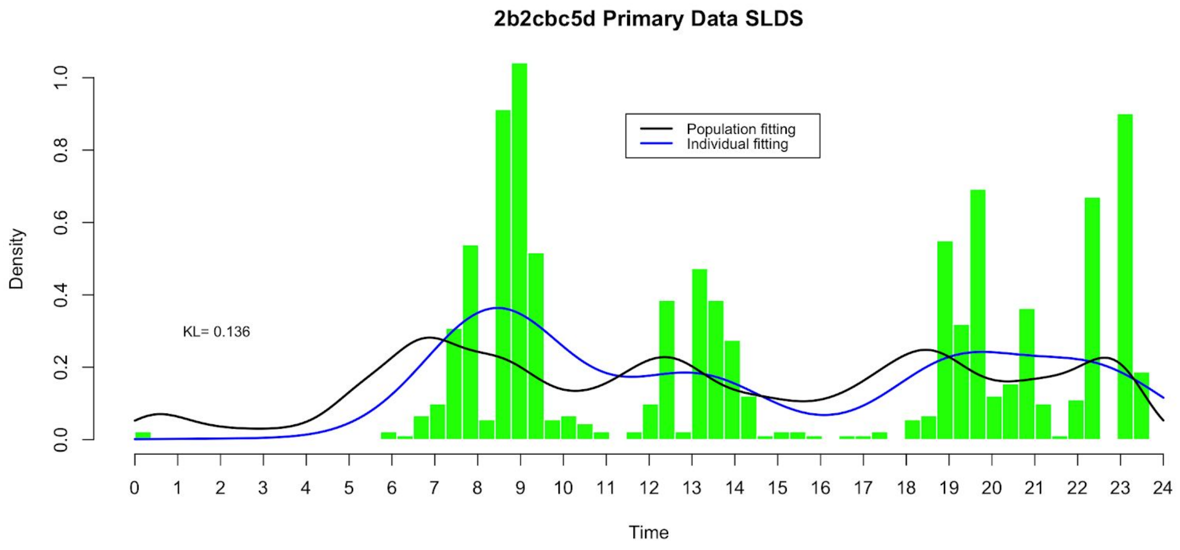
2b2cbc5d Primary Data SLDS
Population fitting
Individual fitting
KL= 0.136
Density
0.0
0.2
0.4
0.6
0.8
1.0
0 1 2 3 4 5 6 7 8 9 10 11 12 13 14 15 16 17 18 19 20 21 22 23 24
Time

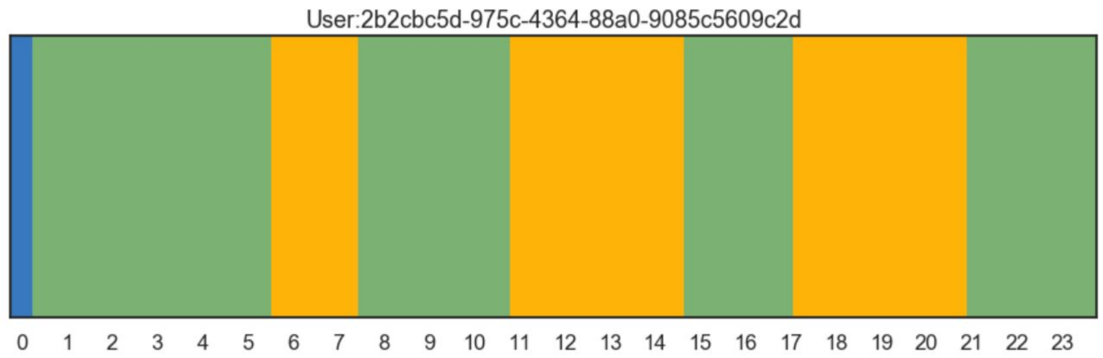
User:2b2cbc5d-975c-4364-88a0-9085c5609c2d
0 1 2 3 4 5 6 7 8 9 10 11 12 13 14 15 16 17 18 19 20 21 22 23

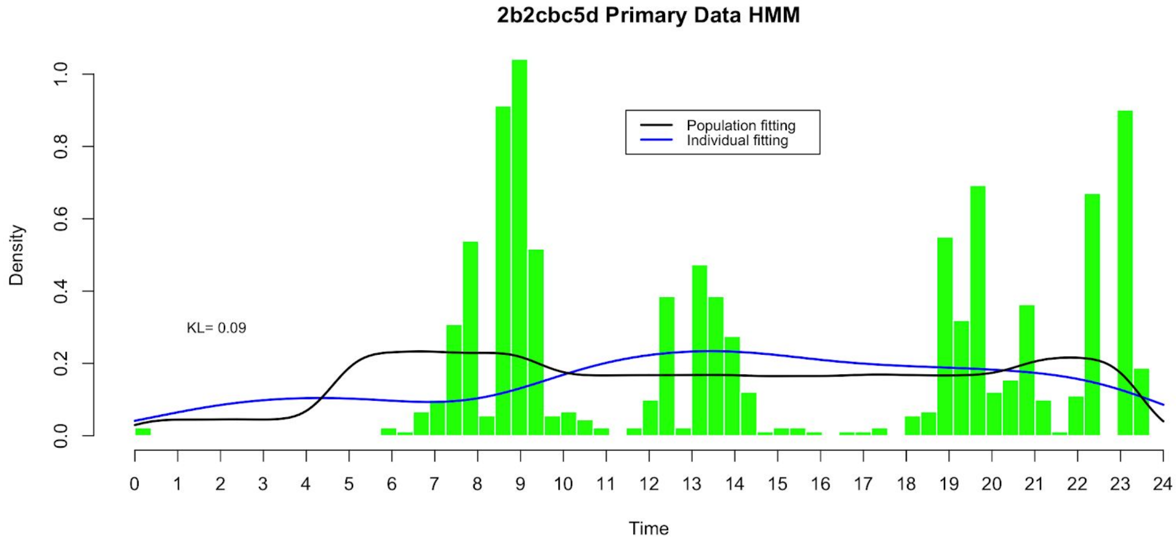
2b2cbc5d Primary Data HMM
Population fitting
Individual fitting
KL= 0.09
Density
Time

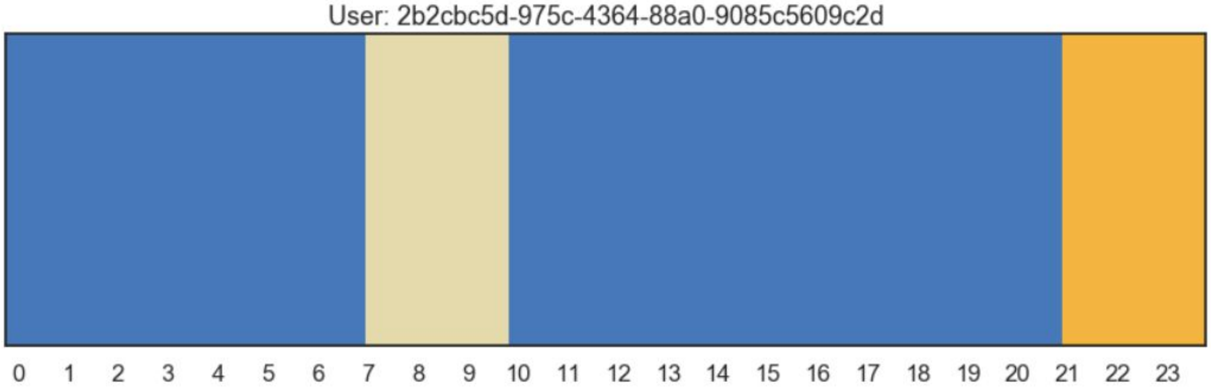
User: 2b2cbc5d-975c-4364-88a0-9085c5609c2d

### 7d439102 Quieter in the middle part of the day

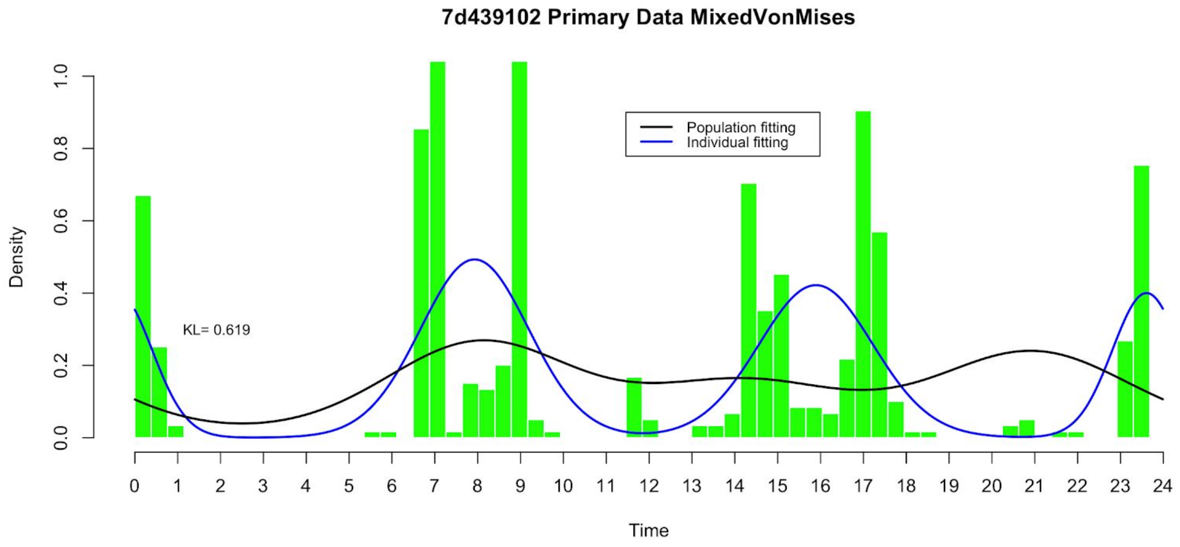


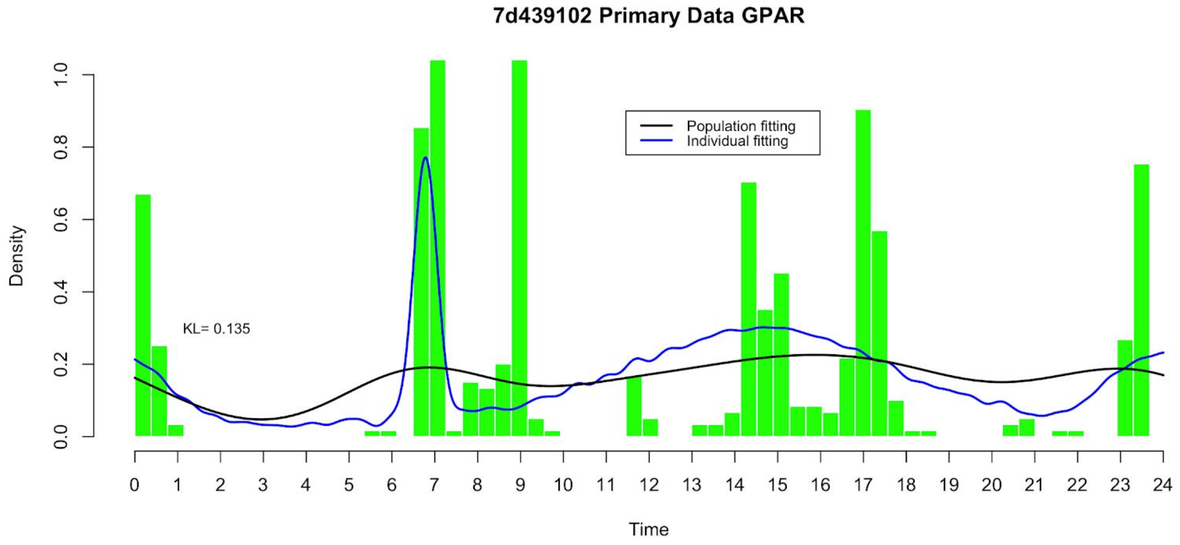

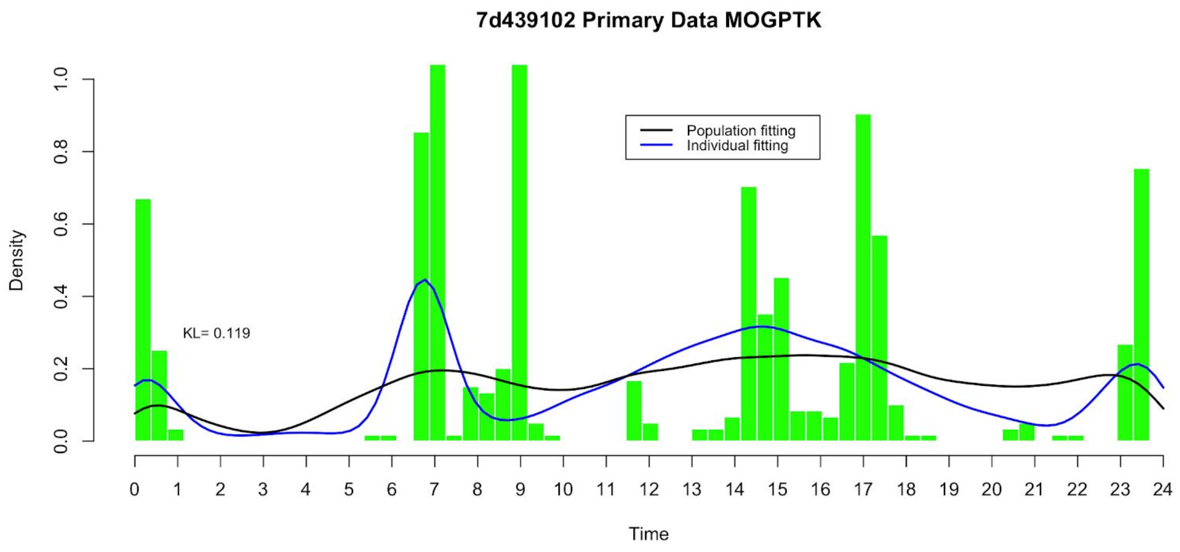
7d439102 Primary Data MOGPTK
Population fitting
Individual fitting
KL= 0.119
Density
Time
0.0
0.2
0.4
0.6
0.8
1.0
0 1 2 3 4 5 6 7 8 9 10 11 12 13 14 15 16 17 18 19 20 21 22 23 24

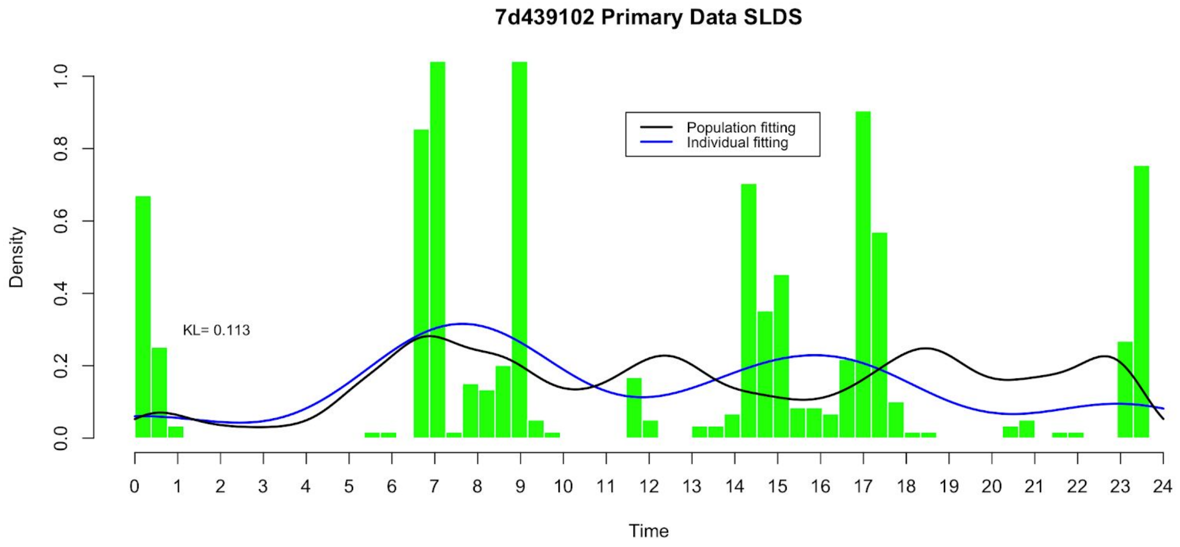
7d439102 Primary Data SLDS
Population fitting
Individual fitting
KL= 0.113
Density
Time
0.0
0.2
0.4
0.6
0.8
1.0
0 1 2 3 4 5 6 7 8 9 10 11 12 13 14 15 16 17 18 19 20 21 22 23 24

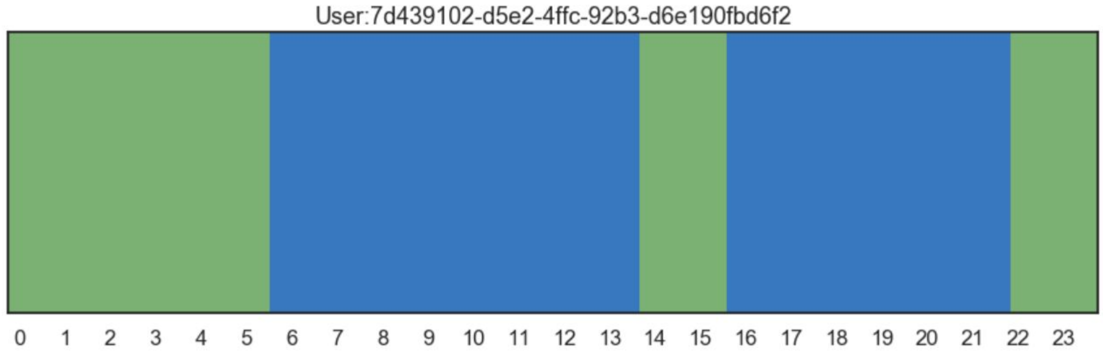
User:7d439102-d5e2-4ffc-92b3-d6e190fbd6f2
0 1 2 3 4 5 6 7 8 9 10 11 12 13 14 15 16 17 18 19 20 21 22 23

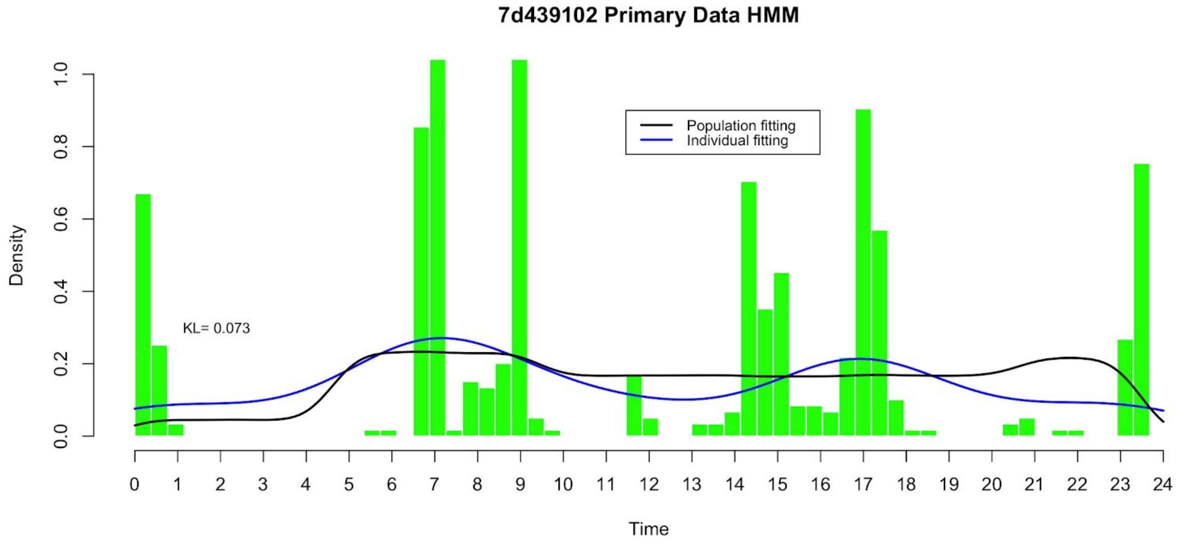

7d439102 Primary Data HMM
Population fitting
Individual fitting
KL= 0.073
Density
Time


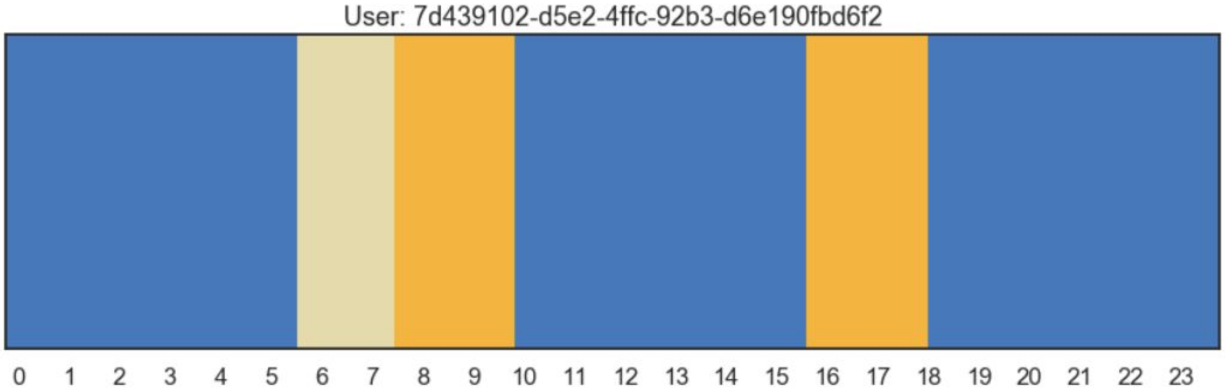

User: 7d439102-d5e2-4ffc-92b3-d6e190fbd6f2

### Population fitting on primary data

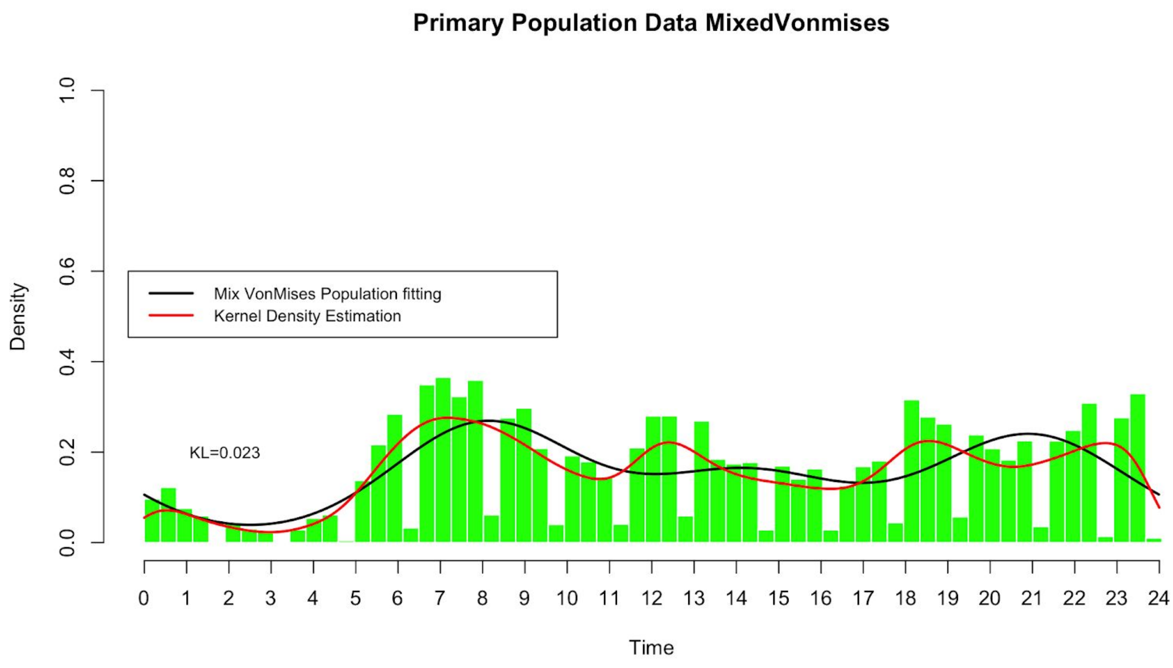


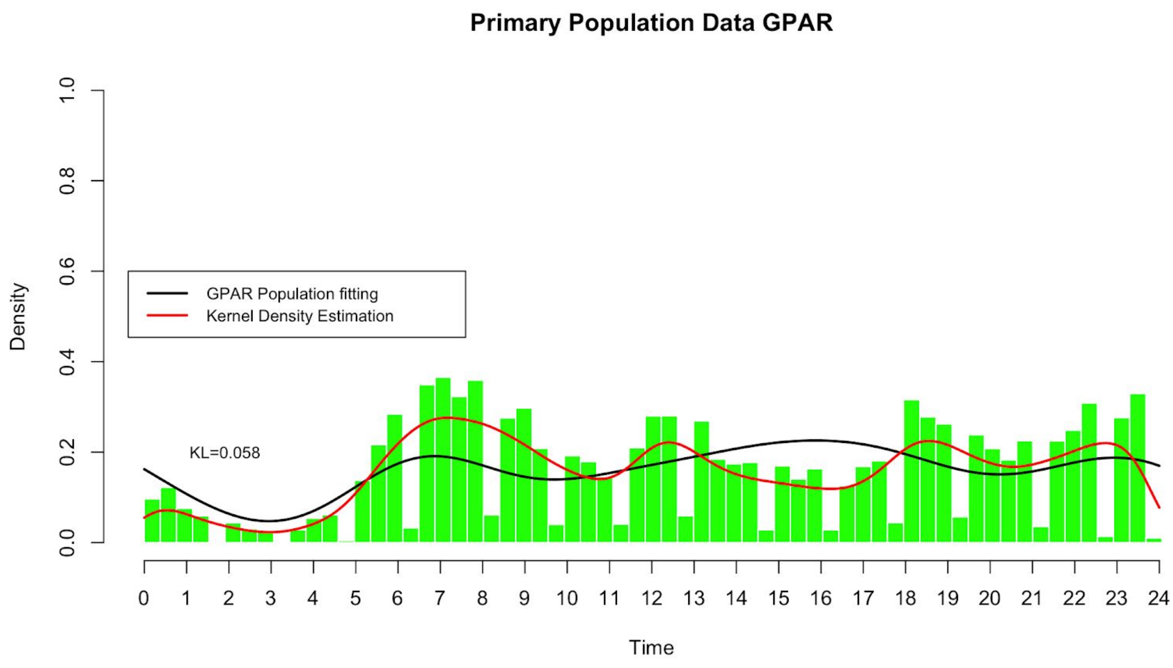

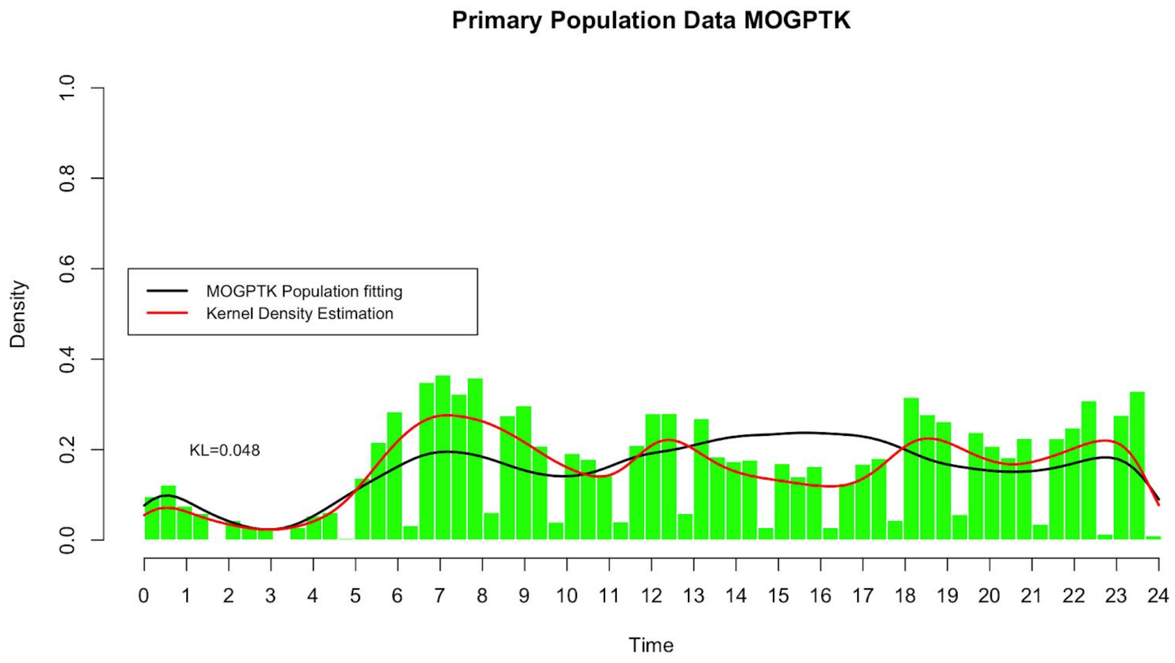
Primary Population Data MOGPTK
Density
Time
MOGPTK Population fitting
Kernel Density Estimation
KL=0.048
0.0
0.2
0.4
0.6
0.8
1.0
0 1 2 3 4 5 6 7 8 9 10 11 12 13 14 15 16 17 18 19 20 21 22 23 24

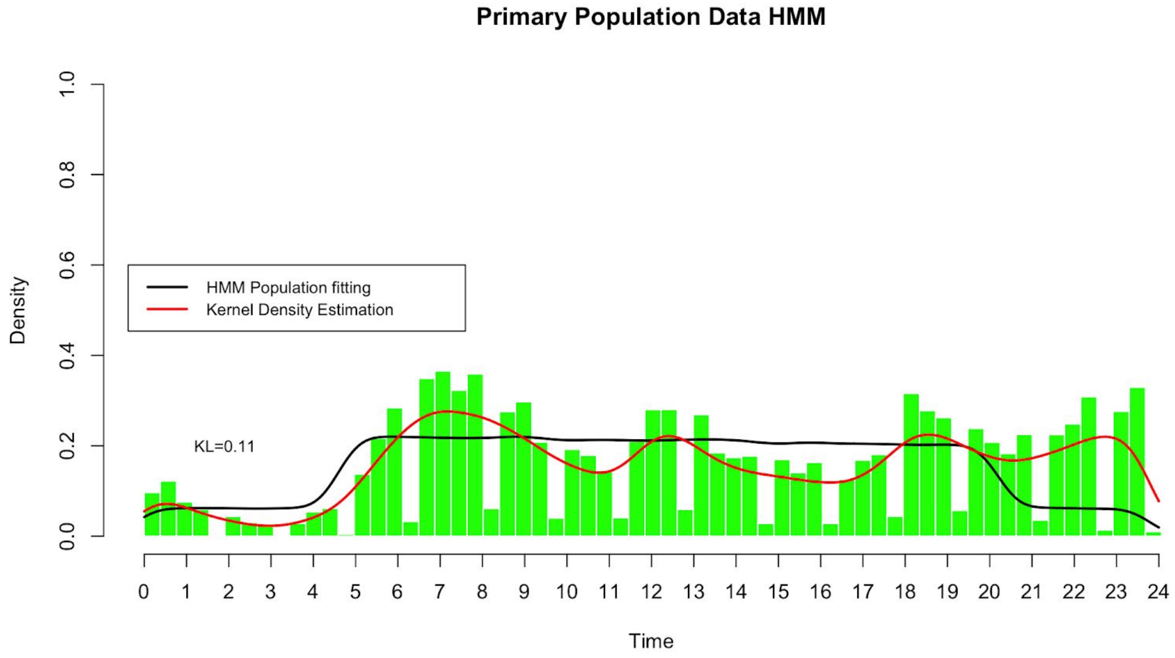
Primary Population Data HMM
Density
Time
HMM Population fitting
Kernel Density Estimation
KL=0.11
0.0
0.2
0.4
0.6
0.8
1.0
0 1 2 3 4 5 6 7 8 9 10 11 12 13 14 15 16 17 18 19 20 21 22 23 24

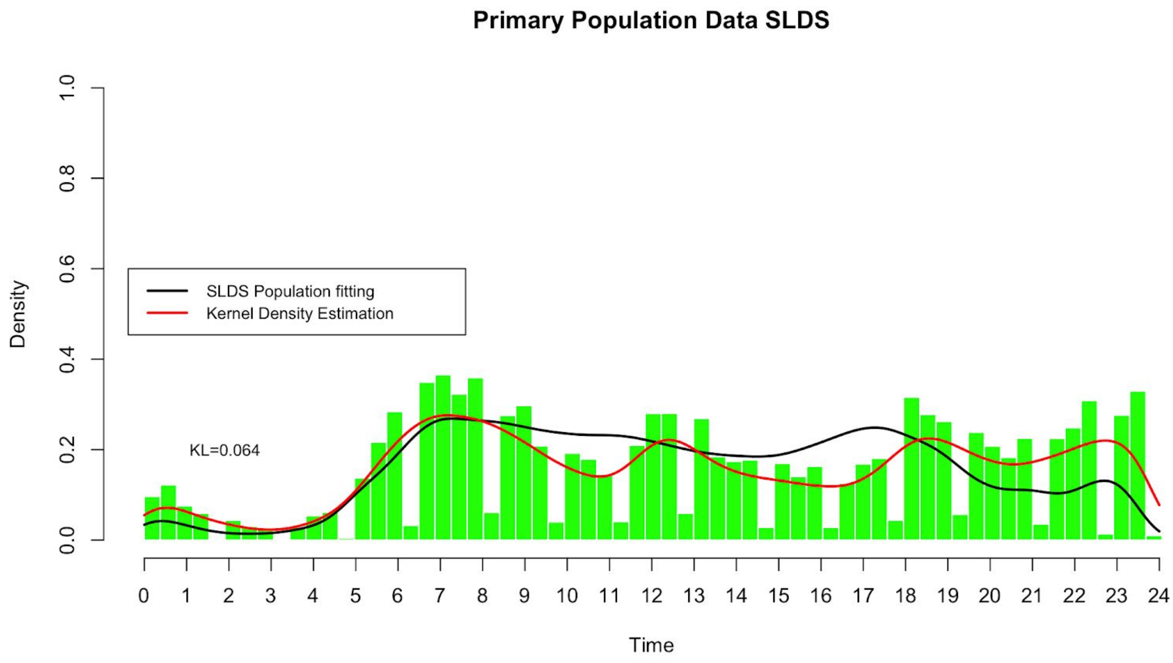
Primary Population Data SLDS
Density
Time
SLDS Population fitting
Kernel Density Estimation
KL=0.064
0.0
0.2
0.4
0.6
0.8
1.0
0 1 2 3 4 5 6 7 8 9 10 11 12 13 14 15 16 17 18 19 20 21 22 23 24
